# Selective and reversible disruption of mitochondrial inner membrane protein complexes by lipophilic cations

**DOI:** 10.1101/2021.03.09.434520

**Authors:** Anezka Kafkova, Lisa Tilokani, Filip Trčka, Veronika Šrámková, Marie Vancová, Tomáš Bílý, Jana Nebesářová, Julien Prudent, Jan Trnka

## Abstract

Triphenylphosphonium (TPP) derivatives are commonly used to target chemical into mitochondria. We show that alkyl-TPP cause reversible, dose- and hydrophobicity-dependent alterations of mitochondrial morphology and function and a selective decrease of mitochondrial inner membrane proteins including subunits of the respiratory chain complexes, as well as components of the mitochondrial calcium uniporter complex. The treatment with alkyl-TPP resulted in the cleavage of the pro-fusion and cristae organisation regulator Optic atrophy-1. The structural and functional effects of alkyl-TPP were found to be reversible and not merely due to loss of membrane potential. A similar effect was observed with the mitochondria-targeted antioxidant MitoQ.

## INTRODUCTION

Mitochondria are organelles involved in a range of functions critical for cell survival^1^. Their functional versatility is accompanied by the remodelling of their network, referred to as mitochondrial dynamics^2^. These dynamic processes encompass continuous cycles of fission and fusion events, which is crucial to answer cellular needs and to adapt to the metabolic state.

Mitochondria are delimited by two membranes, a relatively permeable outer mitochondrial membrane (OMM) and a highly invaginated and selectively permeable inner mitochondrial membrane (IMM). The IMM is highly folded into cristae and contains a range of protein complexes, including those regulating cristae shape, but also respiratory chain complexes, ATP synthase or complexes involved in the regulation of calcium homeostasis (mitochondrial calcium uniporter complex, MCUC, or mitochondrial sodium calcium exchanger, NCLX)^34^. Mitochondrial respiratory chain complexes are responsible for electron transport and the generation of a proton gradient across the IMM, which is used by ATP synthase to produce ATP^5^.

Mitochondria play an important role in many human diseases and being able to manipulate directly their function through targeted drug delivery may be of a substantial benefit^1^. Delivering drugs specifically into mitochondria is challenging because of the double lipid bilayers and the selective permeability of the IMM^6^. However, several strategies for targeting small molecule drugs directly to the mitochondria have been successfully developed. One of the most versatile approaches is to conjugate a small bioactive compound to lipophilic cations such as triphenyl phosphonium (TPP)^7^. These were originally used for measuring of the membrane potential but they have evolved as a targeting strategy for drug delivery^8^.

The chemical properties of the positively charged TPP molecule, combined with the large negative membrane potential in the mitochondrial matrix, enables its free movement through the phospholipid bilayer without relying on a specific uptake mechanism. The three hydrophobic phenyl groups on the TPP moiety shield the positive charge on the phosphorus atom from water molecules to prevent solvation, which lowers the energy preventing the partitioning of the cation into the hydrophobic core of the membrane. At the same time, since the membrane potential across the IMM is up to three times higher compared to the cytoplasmic membrane potential, TPP-conjugated molecules accumulate according to the Nernst equation inside mitochondria^9^. The uptake of the more hydrophobic TPP cations, such as hexyl or decyl-TPP, is more extensive and faster compared to methyltriphenylphosphonium (TPMP), in which the TPP cation is substituted only with a methyl group^10^.

In comparison to unconjugated molecules, multiple studies have shown an increased efficacy of drugs conjugated to TPP^11,12,13,14,15,16^, as well as their potential side effects have been investigated^17,18,19^. MitoQ, a known antioxidant conjugated to decyl-TPP, is already undergoing clinical trials and is close to clinical practice^20^. However, the consequences of TPP mitochondria-targeting moiety itself on different basic mitochondrial behaviour are currently not well characterised. Although the TPP moiety is often seen as inert, several studies have described its significant biological effects, particularly in the inhibition of respiratory chain complexes^19,21^, enzymes of the Krebs cycle,^22^ or the NCLX^18^. The underlying mechanisms of these effects as well as other possible effects on mitochondrial morphology and IMM architecture are still largely unknown.

This study investigates the effect of the alkyl substituted TPP moiety on mitochondrial morphology as well as the stability and turnover of different mitochondrial protein complexes, their impact on mitochondrial bioenergetics and reversibility of these processes. We show that TPP treatment for 24 hours at 2 μM leads to a significant and selective decreased protein levels of different complexes localized at the IMM including complexes of the respiratory chain (RC) and the MCUC. These reversible effects on IMM complexes were accompanied by a decrease of mitochondrial membrane potential and defects in mitochondrial bioenergetics. Finally, confocal and cryo-electron microscopy analysis reveal that TPP treatment leads to a dramatic remodelling of the mitochondrial network characterised by not only mitochondrial fragmentation and swelling, but also by IMM structure alterations.

## METHODS

### Materials

Propyltriphenylphosphonium bromide (131563) and methyltriphenylphosphonium bromide (130079) were purchased from Sigma-Aldrich. Hexyltriphenylphoshonium bromide was obtained from Merck (842007). Actinomycin D (A1410), FCCP (C2920), oligomycin (75351), rotenone (R8875) and antimycin A (A8674) were obtained from Sigma-Aldrich, digitonin from Calbiochem (300410/ D44495). MitoQ and TPP-hexanol were kindly provided by Prof. Michael P. Murphy’s lab (MRC Mitochondrial Biology Unit, Cambridge, UK).

### Antibodies

Rabbit monoclonal anti-TOM20 (ab232589), mouse monoclonal anti-MTCO1 (ab14705), anti-ATP5a (ab14748), anti-VDAC1 (ab14734), anti-SDHB (ab14714), anti-NDFUB8 (ab110242), anti-ubiquinol-cytochromec reductase core protein I (ab110252), anti-GRP75 (ab2799), anti-MIC60 (ab110329), anti-cytochrome c (ab110325), rabbit polyclonal anti-citrate synthase (ab96600) and anti-MICU2 (ab101465) were purchased from Abcam. Antibodies mouse monoclonal anti-Hsp60 (sc136291) and anti-alpha-tubulin (sc23948) were obtained from Santa-Cruz Biotechnology. Antibodies rabbit polyclonal anti-PARP (#9542), anti-cleaved caspase 3 (#9661) and rabbit monoclonal anti-MICU1 (#12524) were purchased from Cell Signalling Technology. Antibodies mouse monoclonal anti-vinculin (V4505), anti-OPA1 (612607), rabbit polyclonal anti-MIC19 (25625-1-AP) and rabbit monoclonal anti-MCU (14997) were purchased from Sigma-Aldrich, BD Transduction Laboratories Proteintech and BioTech, respectively.

### Cell lines, culture conditions and transfections

C2C12 mouse myoblasts cells, and HepG2 human liver cancer cells were obtained from American Type Culture Collection (ATCC). Cells were cultured in low-glucose (1 g/l) Dulbecco’s Modified Eagle’s medium (DMEM) medium supplemented with 10 % fetal bovine serum (FBS) and 0.5 mM L-glutamine in humidified atmosphere at 37°C and 5 % CO2 (all from Gibco-Life Technologies).

TPP compounds were dissolved in ethanol, actinomycin D, FCCP, oligomycin, rotenone and antimycin A were dissolved in DMSO, final concentration of vehicle used for any experiment was below 0.5 %.

For small interfering (si) RNA experiments, C2C12 were transfected using Lipofectamine RNAimax (Invitrogen) with 20 nM siRNA for 3 days, then the transfection was repeated again after 3 days, following manufacturer’s instructions. Non-targeting siRNA was purchased from Invitrogen (4390844) and siRNA targeting OMA1 from Dharmacon (ON-TARGETplus, SMARTpool, L-051170-01-0005).

### Denaturing and native electrophoresis, immunoblot, and immunodetection

Samples for blue-native gel electrophoresis (BNGE) were prepared from digitonised cellular extracts as described previously^23^. For the solubilisation, 2 % digitonin was used. Samples were run through precast Native PAGE Novex 3-12 % Bis-Tris Protein Gels (Thermo Fisher Scientific) according to the manufacturer’s instructions.

Samples for immunoblot were extracted in 1.5 % DDM lysis buffer (D4641, Sigma-Aldrich) or RIPA lysis buffer (R0278, Sigma-Aldrich). 10 μg aliquots of total protein were transferred to hand-cast polyacrylamide gel or to Novex NuPAGE 4-12 % Bis-Tris Precast Gels (Thermo Fisher Scientific).

Proteins were then wet-transferred to PVDF or nitrocellulose membranes, blocked in 10 % non-fat milk (Marvel) in 0.1 % Phosphate buffer saline (PBS)-Tween 20 and incubated with specific primary antibodies overnight. The following day the membranes were washed in 0.1 % PBS-Tween 20, incubated with Goat anti-Mouse IgG (W402B, Promega) or Goat anti-Rabbit IgG (W401B, Promega) secondary antibodies for 1 hour (h), and washed again in 0.1 % PBS-Tween 20. Immunoreactive bands were visualized with ECL Western Blotting Detection Reagents (ThermoFisher) or Amersham ECL Prime Western Blotting Detection Reagent (GE Healthcare) and revealed manually on X-Ray films (Fuji; FOMA LP-T and FOMAFIX) or using a digital Amersham Imager 680 (GE Healthcare).

### Mitochondrial protein solubilisation

50,000 C2C12 cells were seeded in a 6-well plate and cultured overnight. Cells were treated with 2 μM hexyl-TPP or vehicle for 24 h before harvesting. Equal numbers of cells were pelleted and incubated in a mild lysis buffer (1 % Triton-X, 150 mM KCl, 30 mM Tris pH 7.5) for 10 minutes (min) at room temperature (RT), followed by centrifugation at 8,000 x *g* for 5 min at 4 °C. Supernatants with soluble protein fractions were kept and pellets were washed once in PBS and centrifuged at 8,000 x *g* for 5 min at 4 °C. Pellets were resolubilized by vortexing in 2x Laemmli sample buffer and homogenized by passing through silica-based spin column. Soluble and insoluble fractions were analysed for the presence of selected proteins by immunoblot analysis.

### Gene expression analysis

Total RNA extraction was performed using RNeasy Mini Kit (Qiagen, Germany) according to manufacturer’s instructions. RNA concentration was measured using Nanodrop1000 (Thermo Fisher Scientific, USA). DNAse I (Thermo Fisher Scientific) treatment was applied to remove genomic DNA. Four to nine hundred nanograms of total RNA were reverse transcribed using a high-capacity cDNA reverse transcription kit (Applied Biosystems). qPCR was carried out using PowerUp SYBR Green Master Mix (Thermo Fisher Scientific) and gene-specific primers: mtco1: 5-gacttgcaaccctacacgga-3, 5- gatggcgaagtgggcttttg-3; actb: 5-cctgaaccctaaggccaacc-3, 5-atggcgtgagggagagcatag-3; coxIV: 5-gctccgtcttggtcttccg-3, 5-cttcacaacactcccatgtgct-3; nd6: 5-tattgccgctaccccaatcc-3, 5-gggggatgttggttgtgtttg-3; ndufb8: 5-tggggtgaaccgatacactg-3, 5-agcacgtagagggaaggaca-3, or Fast Advanced Master Mix and TaqMan primers and probes (oma1-Mm01260328_m1, csnk2a2 - Mm01260328_m1, Thermo Fisher Scientific). Data were analysed using the comparative ΔΔCt method with values normalized to β-actin or Csnk2a2 levels. Specificity of PCR reaction for each amplicon in Sybr Green reaction was tested by melting curve analysis.

### Flow cytometry

Changes in the mitochondrial membrane potential were determined as the changes in tetramethylrhodamine methyl ester (TMRM, T668, Thermo Fisher Scientific) fluorescence according to the manufacturer’s instructions with few modifications. Briefly, cells were treated with 2 μM hexyl-TPP, 2 μM FCCP or vehicle for 24 h. Then, cells were washed with PBS and incubated in staining solution (50 nM TMRM in Hank’s Balanced Salt Solution with 2 μM vehicle/hexyl-TPP/FCCP) at 37 °C for 30 min. Finally, cells were harvested, centrifuged at 350 x *g* for 3 min and resuspended in Hank’s Balanced Salt Solution with 3% FBS. Cells were then analysed using a BD Verse flow cytometer. 10,000 events per sample were used and TMRM fluorescence was measured using the PE channel, with TMRM excitation at 488 nm and emission at 586 nm.

### Immunofluorescence

10,000 C2C12 cells were seeded on a glass coverslip in 24-well plates and incubated overnight. Cells were then treated with 2 μM hexyl-TPP for different timepoints. The following steps were performed as previously described^24^. Briefly, the medium was removed, and cells were fixed with 5 % paraformaldehyde at 37°C for 15 min, washed in PBS, followed by quenching with 50 mM ammonium chloride in PBS, washed with PBS and permeabilized with 0.1 % Triton X-100 in PBS for 10 min. Next, cells were washed with PBS and blocked with 10 % FBS in PBS. Cells were then incubated with the primary antibody (anti-TOM20) in 5 % FBS in PBS for 2 h at RT. After three washes with 5 % FBS in PBS, cells were incubated with secondary antibodies (Alexa Fluor 594, A11012, Thermofisher Scientific) in 5 % FBS in PBS for 1 h RT, washed three times with PBS and mounted onto slides using fluorescence mounting medium (Dako).

### Confocal imaging

Fixed cells were imaged using 100X objective lens (NA1.4) on a Nikon Eclipse TiE inverted microscope with appropriate lasers using an Andor Dragonfly 500 confocal spinning disk system, equipped with a Zyla 4.2 PLUS sCMOS camera coupled with Fusion software (Andor). 7 stacks of 0.2 μm each were acquired. Images were compiled by “max projection” using Fiji software^25^ and cells were classified as fragmented, tubular and hyperfused, similarly as previously described.^26^ In addition, we included aggregated mitochondria as a category of the mitochondrial morphology phenotype. For mitochondrial area and number analysis, 225 μm^2^ of region of interest (ROI) was selected from max projection images and further analysed using the “Analyse particles” plugin in Fiji with a minimum area of 0.1 μm^2^ as described previously^26^. Three independent experiments were performed. The numbers of cells quantified per conditions are indicated in figure legends, specifically for each experiment.

### Transmission electron microscopy

#### High-pressure freezing/freeze substitution (HPF/FS)

After 24 h of treatment with 2 μM hexyl-TPP or vehicle, C2C12 cells were gently scraped and centrifuged at 350 x *g* for 5 min. Cells were frozen using a Leica EM ICE high-pressure freezer in the presence of 20 % bovine serum albumin (w/v). Frozen cells were cryo-substituted in medium containing 1 % osmium tetroxide in 100 % acetone at −90 °C for 96 h. Then, temperature was raised (5 °C/14 h) to −20°C and after 24 h to 4 °C (3 °C/18 h). At RT, specimens were washed three times in 100 % acetone and infiltrated with 25 %, 50 %, 75 % solutions of Embed 812 resin (EMS) in acetone for 1 h. Specimens were incubated in 100 % resin overnight and finally embedded and polymerized at 60 °C for 48 h. Ultrathin sections were counterstained in saturated ethanolic uranyl acetate and lead citrate and observed using TEM JEOL 1010 working at 80 kV.

#### Tokuyasu cryosectioning

After 24 h of treatment with 2 μM hexyl-TPP or vehicle, C2C12 were fixed in 4 % formaldehyde with 2.5 % glutaraldehyde in 0.1 M HEPES. After 1 h incubation at RT and at 4 °C overnight, cells were washed with 0.1 M HEPES and cryoprotected in 2.3 M sucrose for 72 h at 4 °C and frozen by plunging into liquid nitrogen. Ultrathin sections (100 nm) were cut at −100 °C, picked-up with 1.15M sucrose/1 % methylcellulose solution (25 cp, Sigma). Sections were incubated with gold nanoparticles (15 nm) for 3 min, washed in dH2O, and contrasted/embedded using a mixture of 2 % methylcellulose and 3 % aq. uranyl acetate solution (9:1) for 10 min. Samples were observed with a MegaView III camera mounted on a JEOL 1010 TEM. Electron tomography was done on a 200 kV TEM JEOL 2100F equipped with motorized tilt stage and a Gatan K2-summit camera.

#### Electron tomography

SerialEM^27^ was used to automated acquisition of tilt series according to Saxton^28^ scheme in range ± 70° with 131 projections. The tomogram reconstruction and model creation were done in ETomo software package^29^. Reconstructed tomograms with isotropic pixel size 0.56 nm were filtered using low pass Fourier filter in Fiji software package^25^.

### Oxygen consumption measurement

C2C12 cells treated with vehicle or 2 μM hexyl-TPP were seeded overnight in 24-well culture plate at 30,000 cells/well. Next day, culture medium was removed and replaced with XF Assay Media from Seahorse XF DMEM medium supplemented with 5.5 mM glucose, 1 mM pyruvate, 4 mM glutamate and 2 μM hexyl-TPP/vehicle and incubated at 37°C for 1 h. Oxygen consumption rate (OCR) was measured using the Seahorse XF24 Extracellular Analyzer. The assay consisted of three basal rate measurements, followed by sequential injections of 1 μM oligomycin, 2 μM FCCP and 1 μM rotenone + 1 μM antimycin. Three rate measurements were performed after each injection. OCR was normalized to protein by the Bradford assay. The results are expressed as the average of three independent experiments.

### Statistics

Graphs show mean ± standard error of mean (SEM) from three independent experiments. Statistical significance was assessed using one-way ANOVA or two-way ANOVA with Tukey’s multiple comparison or unpaired t-test in GraphPad Prism software (GraphPad Prism 8, https://www.graphpad.com), *p < 0.05, **p < 0.01, ***p < 0.001.

## DATA AVAILABILITY STATMENT

All data are provided in the main article and in the supplementary. Raw data from the current study are available from the corresponding author on request.

## RESULTS

### 1. TPP derivatives affect OXPHOS complex protein levels in a dose-, time- and hydrophobicity-dependent manner

Previous studies have shown that different alkylated TPP cations inhibit cellular respiration after only a few minutes of exposure^19,21^. These studies were exclusively focused on functional analyses and did not provide any evidence of possible structural defects of the mitochondrial respiratory chain induced by the TPP molecules or the potential effects of longer-term treatment. To address these issues, we exposed C2C12 cells to three alkyl-TPP derivatives with an increasing alkyl chain length and increasing hydrophobicity, namely methyltriphenylphosphonium (TPMP), propyltriphenylphosphonium (propyl-TPP or 3-TPP) and hexyltriphenylphoshphonium (hexyl-TPP or 6TPP) (Fig. S1A-C), for 24 hours and analysed the effect on steady state protein levels of components of OXPHOS complexes including MTCO1 for cytochrome c oxidase (Complex IV, CIV) and SDHB for succinate dehydrogenase (Complex II, CII) (Figure 1A-C, and Fig. S2A-C, *S10*).

**Figure 1.**
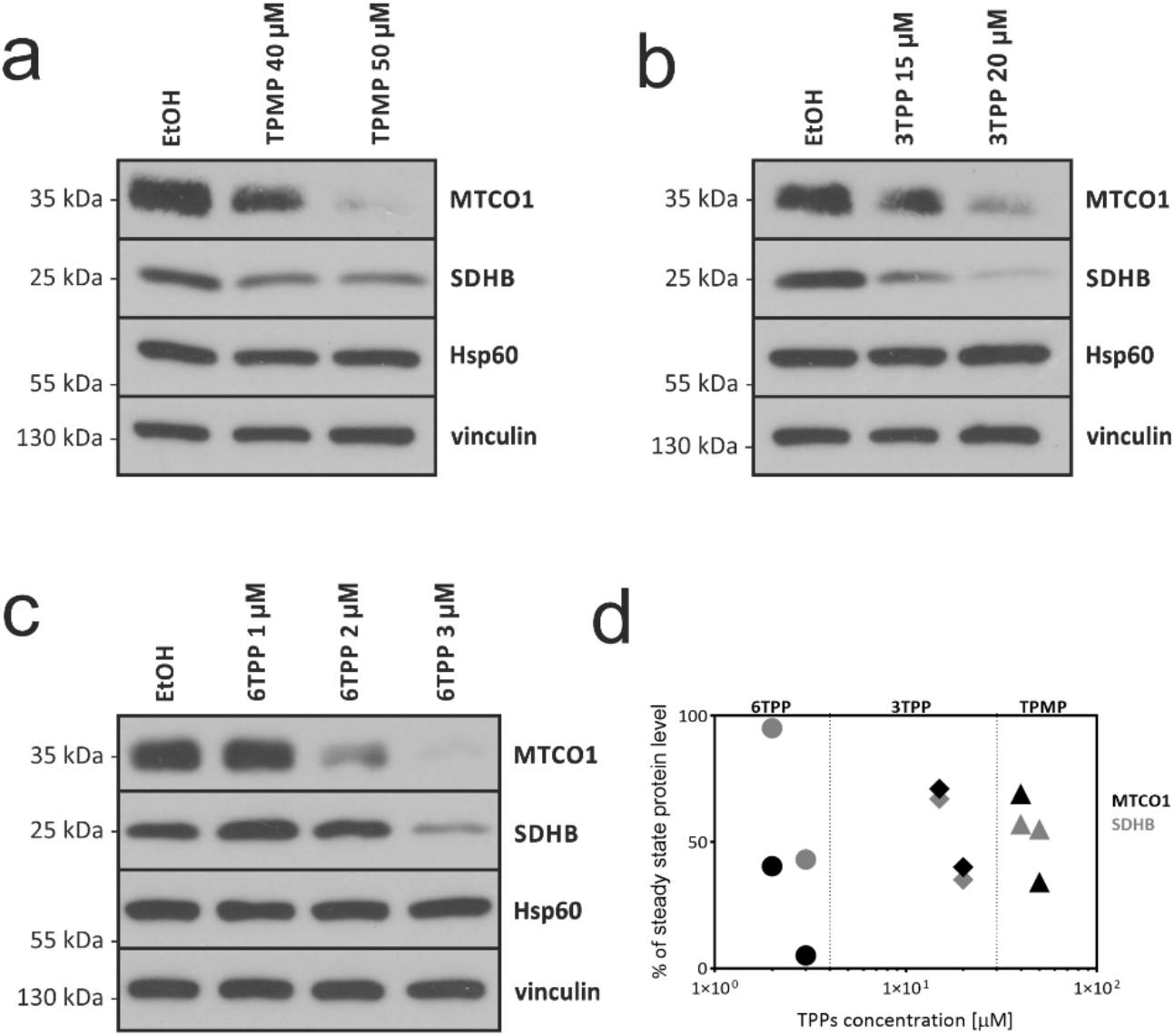
Treatment with TPP derivatives leads to a decrease in selected subunits of OXPHOS complexes. Representative immunoblot for steady-state levels of respiratory chain complex IV (anti-MTCO1) and complex II (anti-SDHB) from C2C12 cells treated for 24 hours with ethanol (EtOH) or different concentrations of methyltriphenylphosphonium (TPMP) (**A**), propyl-TPP (3-TPP) (**B**), and hexyl-TPP (6-TPP) (**C**). Vinculin and Hsp60 were used as loading controls. **D.** Graph showing the reduction in steady state protein levels upon exposure to TPPs with different hydrophobicity (compared to vehicle, x-axis log scale).

Incubation of C2C12 cells with the compounds led to a concentration- and hydrophobicity-dependent decrease in the levels of inner mitochondrial membrane proteins; the subunits of CIV (MTCO1) and CII (SDHB). While hexyl-TPP caused a strong reduction of protein levels at 2 μM, 15 and 40 μM of propyl TPP and TPMP, respectively, had to be used to achieve a similar decrease in the amounts of respiratory chain subunits (Figure 1D). This is in accordance with the literature Ross et al.^10^ where increasing the hydrophobicity of TPP compounds (i.e. the length of alkyl chain) correlates with an increased ability of TPP derivatives to cross the plasma and mitochondrial membranes which facilitate their accumulation in mitochondria. Therefore, we decided to mainly use the most active compound (hexyl-TPP) in further studies.

Next, we analysed the effect of hexyl-TPP treatment on the steady-state protein levels of one representative subunit of each respiratory complex, NADH:ubiquinone oxidoreductase, CI, (NDUFB8), CII (SDHB), cytochrome c – oxidoreductase, CIII (COREI), CIV (MTCO1) as well as of ATP synthase (ATP5a) at different timepoints (Figure 2A, Fig. S3A, *S11A*). While the levels of ATP5a remained unchanged after 24 h of treatment, the levels of other subunits of the RC reduced over time. Of these tested proteins, the most sensitive to hexyl-TPP treatment were MTCO1 and NDUFB8, while SDHB and CORE1 were less affected. The decrease of MTCO1 and NDUFB8 was statistically significant at 6 or 24 hours, respectively; however, there was already a strong decrease at 2 h (MTCO1) and 6h (NDUFB8) of treatment. The change of CORE1 did not reach statistical significance (Fig. S3A).

**Figure 2.**
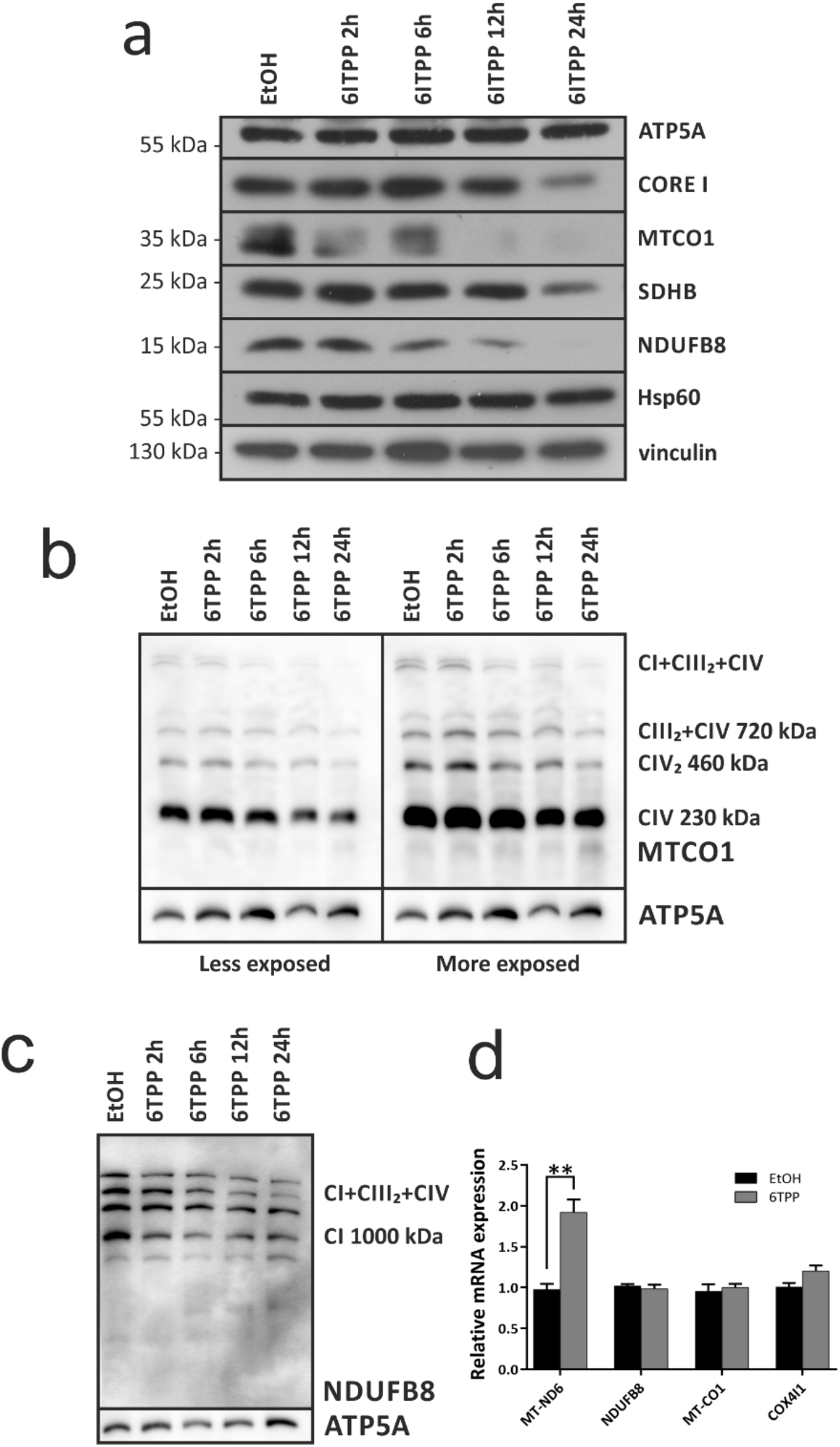
Treatment with hexyl-TPP leads to a decrease of OXPHOS complexes in a time dependent manner. **A.** Steadystate protein levels of respiratory chain components, NDUFB8 of complex I, SDHB of complex II, CORE I of complex III, MTCO1 of complex IV and of complex V (ATP5A) in C2C12 cells treated with ethanol (EtOH) or with 2 μM hexyl-TPP over different timepoints as detected by immunodetection with specific antibodies against the indicated subunits. Vinculin and Hsp60 were used as loading controls. **B, C.** 1D-BNGE and immunodetection analysis showing the decrease of fully assembled (B) complex IV, detected with anti-MTCO1 antibody and (C) complex I, detected with an anti-NDUFB8 antibody, after 2 μM hexyl-TPP treatment at different timepoints, in C2C12 cells. ATP5A was used as loading control. **D**. Quantification of mRNA levels in C2C12 cells treated with ethanol or hexyl-TPP for 24 hours by qRT-PCR normalized to actin mRNA and expressed relative to control cells. Results represent three independent experiments, error bars indicate mean ± SEM, ** indicate p < 0.01 in unpaired student t-test.

To investigate the effect of hexyl-TPP on the assembly of IMM proteins into their native respiratory chain complexes and supercomplexes^30^, we then performed Blue Native PAGE (BN-PAGE) analysis (Figure 2B, C, Fig. *S11 B,C*). C2C12 cells were treated with 2 μM hexyl-TPP at different timepoints, 2, 6, 12 and 24 hours. The effects of these treatments on CIV stability and formation were evaluated by immunodetection with the antibody recognising MTCO1, after the native electrophoresis of samples solubilised with digitonin (Figure 2B). We observed a decrease in the amounts of CIV bound to the supercomplexes after 6 hours of treatment, whereas the decrease of ‘free’ CIV was visible at 12 hours and more pronounced after 24 hours of treatment. The observed early effect on the supercomplexes formation could be due to specific disassembly or lack of assembly of CIV into these structures. Alternatively, it could be a consequence of earlier disruptions to other components of the respiratory chain supercomplexes. Therefore, we further performed BN-PAGE analysis followed by immunoblotting against NDUFB8 and we also observed defects in assembly of CI. Given that NDUFB8 is a CI supernumerary subunit necessary for the assembly of the holocomplex^31^, this can explain why the levels of supercomplexes were already reduced at 6 hours of exposure to hexyl-TPP (Figure 2B, C). Together, these results suggest that the hexyl-TPP treatment may impact newly synthetised or unassembled protein subunits of the RC.

A similar effect on steady-state protein levels was also observed in human HepG2 cells treated with hexyl-TPP, suggesting that this is not a cell-line specific phenomenon (Fig. S3B, *S18A*). To exclude the possibility of a decreased mitochondrial protein solubility induced by hexyl-TPP treatment, we also performed mild cellular lysis followed by SDS extraction of insoluble pelleted proteins (Fig. S3C, *S18B*). As MTCO1 protein remained detergent-soluble after the treatment with hexyl-TPP, we excluded the possibility that the observed decrease was simply due to protein aggregation.

We also tested the hypothesis that TPP treatment may affect the expression of genes coding for different subunits of CI and CIV, whose steady state protein levels were the most affected by hexyl-TPP treatment (Figure 2D). The nuclear encoded mRNA transcript levels of genes for CI (NDUFB8) or CIV (COX4/1) remained unchanged. However, while transcript levels of the mitochondrial encoded gene for CIV (MTCO1) is unchanged, transcript levels of gene for CI (MT-ND6) were increased. Therefore, this experiment ruled out the possibility of hexyl-TPP decreasing mtDNA transcription or downregulation of nuclear genes encoding RC components.

### 2. Hexyl-TPP treatment results in a decrease in MCUC steady state protein levels and enhanced OPA1 cleavage

In order to elucidate whether the decrease in the levels of OXPHOS-related proteins upon exposure to hexyl-TPP was target-specific or due to a global reduction of mitochondrial proteins, the levels of OMM (TOM20, VDAC1), matrix (GRP75 and citrate synthase), IMM (MIC60, MIC19, MCU) and inter-membrane space (IMS) (MICU1, MICU2, cytochrome *c*) proteins were determined using immunoblot analyses (Figure 3A-D, *S12 A-C, S13A*). We observed no changes in the levels of OMM and matrix proteins. Interestingly, whilst the levels of the mitochondrial calcium uniporter (MCU) and its associated gatekeepers (MICU1 and MICU2) were decreased (Fig. S4A), other steady state protein levels of IMM and IMS proteins such as MIC60, MIC19 and cytochrome c, remained unchanged after a 24-hour treatment.

**Figure 3.**
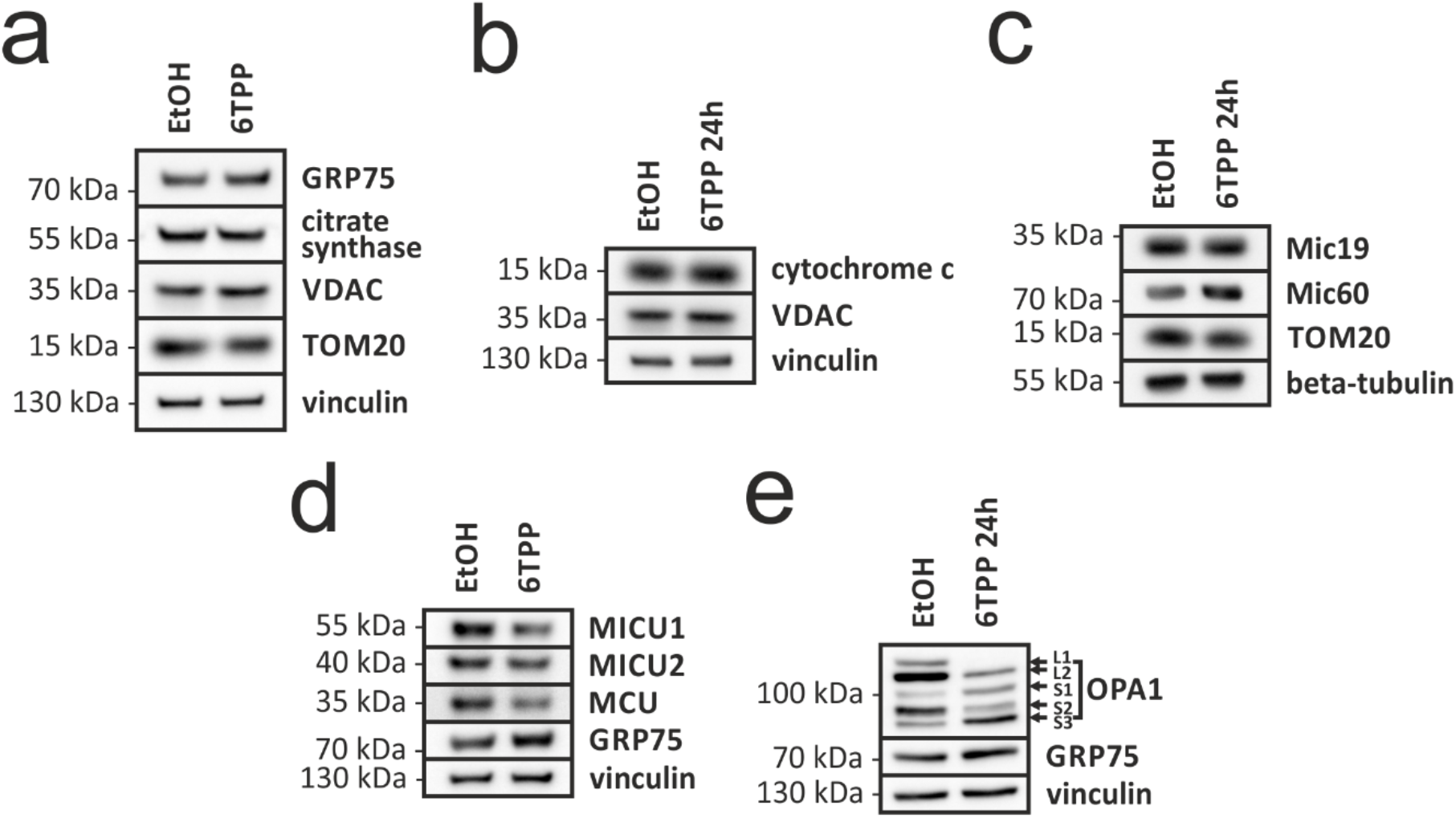
Treatment with hexyl-TPP leads to the decrease of MCUC subunits and OPA1 cleavage. **A**. Steady-state levels of OMM and matrix proteins from C2C12 cells treated with vehicle (EtOH) or 2 μM hexyl-TPP after 24h treatment as detected by immunoblotting. Vinculin was used as a loading control. **B-D.** Steady-state levels of IMM and IMS proteins from C2C12 cells treated with vehicle (EtOH) or 2 μM hexyl-TPP after 24h treatment. Vinculin or beta-tubulin were used as a cytoplasmic loading control and VDAC1, TOM20 or GRP75 were used as a mitochondrial loading control. **E.** The effect of 2 μM hexyl- TPP treatment in C2C12 on OPA1 processing and stability. Vinculin and GRP75 were used as loading controls.

We also analysed the protein level of the IMM GTPase OPA1, which is involved in the regulation of both mitochondrial IMM fusion and cristae maintenance^32^. In response to different stress stimuli including depolarizing agents such as FCCP or others, a transient depolarization of the IMM can lead to the activation of the metalloprotease OMA1, which cleaves the long OPA1 isoforms (L-OPA1) into shorter OPA1 (S-OPA1) isoforms^32,33,34^. We therefore detected OPA1 isoforms after 24 h of 2 μM hexyl-TPP exposure. We observed a reduction in L-OPA1 isoforms (L1 and L2), which was accompanied by the accumulation of the shorter OPA1 isoforms (S1 and S3) resulting from OPA1 cleavage (Figure 3E, *S13B*). OPA1 cleavage was detected also at earlier timepoints similar to the decrease of steady state protein levels of RC complexes (Fig. S5, *S18C*). However, OPA1 cleavage does not appear to precede the loss of OXPHOS subunits, rather it occurs simultaneously. Moreover, we found no inhibition of the observed effect of hexyl-TPP on OXPHOS subunits in cells with a decreased expression of OMA1 after a transfection with OMA1 siRNA despite the fact that the effect of OMA1 siRNA treatment on OPA1 processing was significant and comparable to OMA1 KO cells (Fig. S6)^34^. Both these findings suggest that OMA1 activation and OPA1 cleavage are unlikely to be a causal agent in the cascade of TPP-mediated IMM protein loss but either a concurrent effect of an upstream event or possibly a consequence of IMM protein degradation further.

Collectively these results indicate that rather than a global, non-specific effect on the mitochondrial proteome, the effect of hexyl-TPP is selective for particular mitochondrial proteins and while it is accompanied by a mitochondrial stress leading to OPA1 cleavage, this mechanism is unlikely to explain the observed changes of IMM protein levels, but rather is a downstream effect leading to the observed changes in mitochondrial morphology.

### 3. Hexyl-TPP treatment alters mitochondrial morphology

The cleavage of IMM L-OPA1 and the resulting accumulation of short OPA1 isoforms have been associated to mitochondrial fragmentation as well as disorganised and swollen cristae structure^34^. To investigate whether the accumulation of short OPA1 isoforms after hexyl-TPP exposure results in a similar phenotype, we used confocal and electron microscopy to assess mitochondrial morphology and cristae ultrastructure. We observed that hexyl-TPP treatments leads to a remodelling of the mitochondrial network (Figure 4).

**Figure 4.**
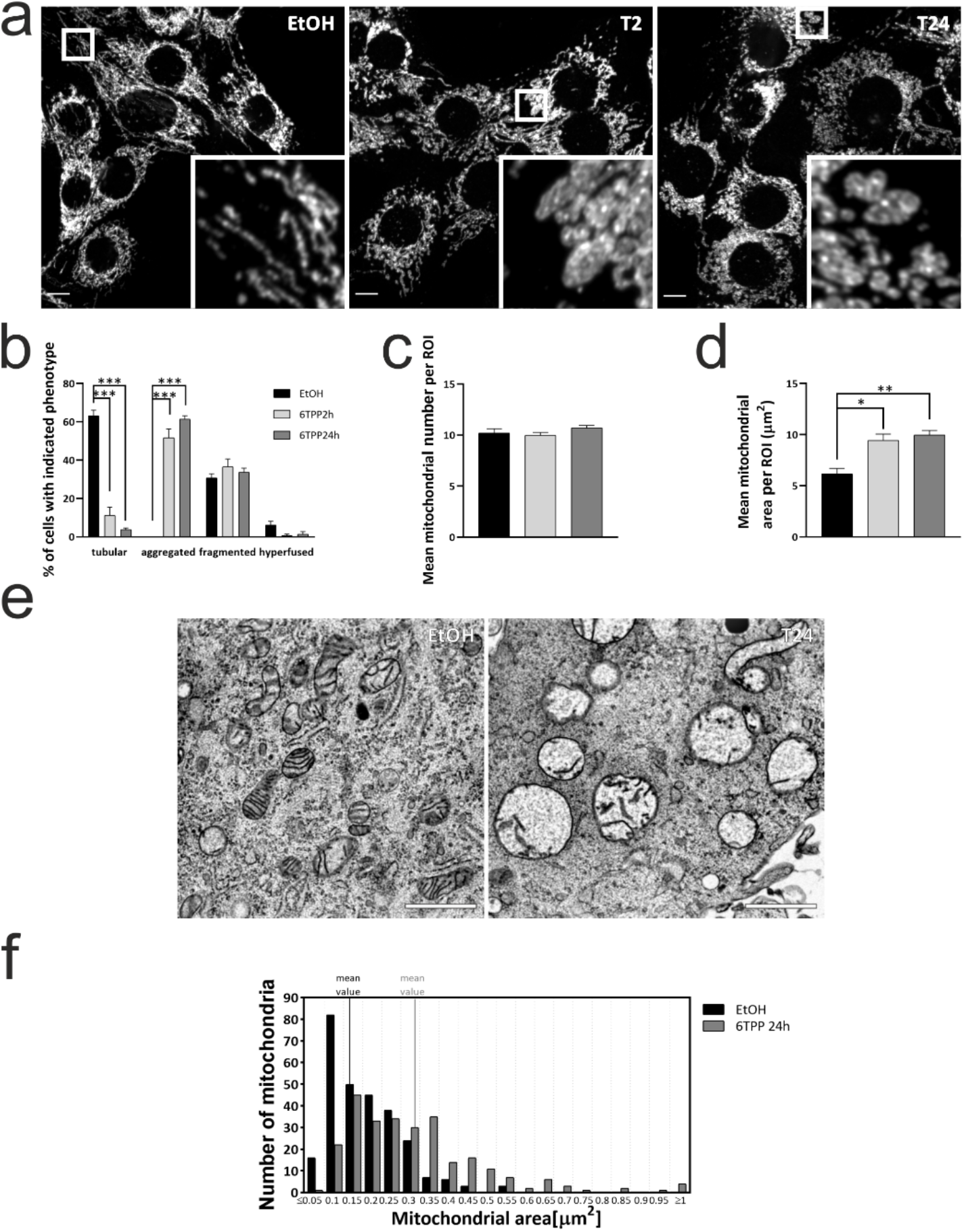
Treatment with hexyl-TPP alters mitochondrial morphology. **A**. Representative confocal images of mitochondrial morphology in C2C12 cells treated with vehicle (EtOH) or 2 μM hexyl-TPP for 2 or 24 hours observed by confocal microscopy. Mitochondria were labelled using an anti-TOM20 antibody. Scale bars, 10 μm. **B.** Quantification of mitochondrial morphology from (A). **C.** Mitochondrial morphology quantified for mean mitochondrial number for each region of interest (ROI). **D.** Mitochondrial morphology quantified for mean mitochondrial area per mitochondrion. All data are shown as mean ± SEM, for B at least 50 cells per condition, for C and D 15 cells for each experiment were quantified; three independent experiments were performed. For B two-way ANOVA and Tukey’s multiple-comparisons test was used. For C and D ordinary one-way ANOVA and Tukey’s multiple comparisons test was used. **E.** Representative images from transmission electron microscopy representative images of mitochondrial ultrastructure of C2C12 cells treated with vehicle (EtOH) or 2 μM hexyl-TPP for 24 hours and prepared by HPF/FS protocol. Scale bars, 0.5 μm. **F.** Quantification of mitochondria area from E. The area of at least 250 mitochondria was quantified for both, control (EtOH) and treatment (6TPP).

After 2- and 24-hours hexyl-TPP treatment, we did not observe increased mitochondrial fragmentation using confocal microscopy, instead mitochondria were aggregated close to the perinuclear area and were swollen with an increase in the mitochondrial area (Figure 4A-D). This phenotype was confirmed by electron microscopy (Figure 4E, F), where we observed not only rounded mitochondria but also an increase of the mitochondrial area. Electron microscopy also revealed a loss of the cristae structure (Figure 4E).

Interestingly, after a shorter one-hour hexyl-TPP treatment we observed a significantly increased number of cells with fragmented mitochondria compared to control. This phenotype was characterised by a reduction in mitochondrial area, which was accompanied by an increase in mitochondrial number per unit area (Fig. S7A-D). These results suggest that short-term hexyl-TPP treatment induced first mitochondrial fragmentation, which shifted towards mitochondrial aggregation and swelling upon longer treatments.

To better characterise the changes in mitochondrial ultrastructure, we further performed a 3D reconstruction of mitochondria in chemically fixed and cryo-sectioned cells. We confirmed the loss of cristae in mitochondria from cells exposed to 2 μM of hexyl-TPP for 24 hours compared to control mitochondria. The negative staining of mitochondrial membrane enabled us to visualize the IMM, which remained unchanged after the treatment with hexyl-TPP and no rupture in the IMM was detected (Figure 5 A-D). This phenotype, i.e. loss of cristae as well as mitochondrial fragmentation and swelling are often associated with cell death^35^. However, immunoblot analysis of caspase 3 and PARP cleavage, two hallmarks of the apoptotic program^36,37,38^, did not suggest apoptosis, even after a prolonged hexyl-TPP exposure (72 h) (Fig. S7E, *S18E*).

**Figure 5.**
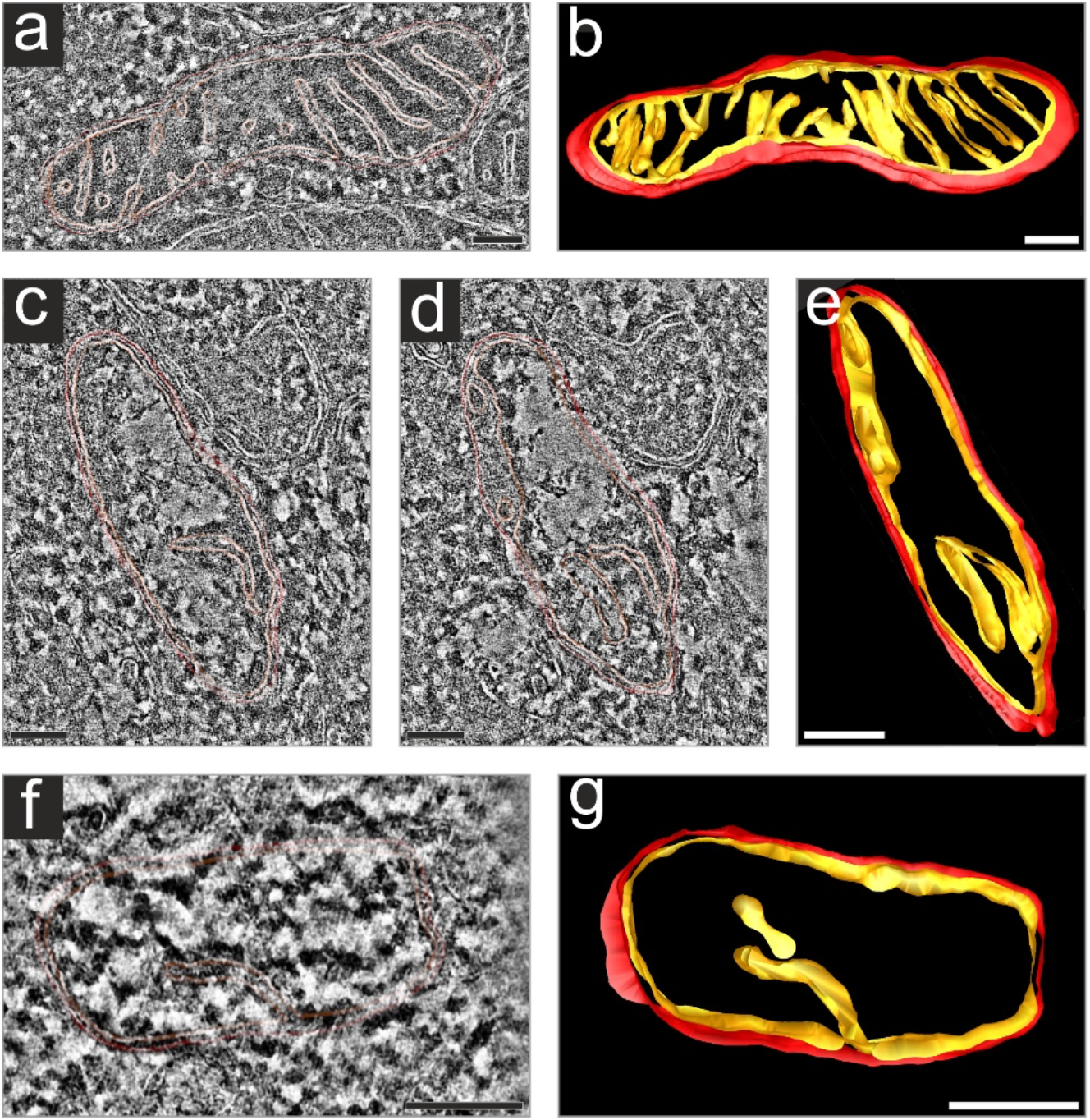
Hexyl-TPP treatment leads to cristae loss but does not impact the inner boundary membrane. Representative images of tomographic slice and 3D reconstruction of mitochondria in vehicle (EtOH) and hexyl-TPP treated cells, mitochondrial membranes are represented in red (outer) and orange (inner). **A, B** control, **C-G** 24 hours exposure to 2 μM hexyl-TPP. The difference in height between C and D is ~10 nm. Each tomogram virtual slice represents thickness 0.56 nm. Scale bars, 100 nm.

### 4. Loss of IMM proteins is membrane potential independent

Acute exposure to TPP derivatives has been reported to decrease the mitochondrial membrane potential^21^. We also showed that chronic treatment (24 hours) with hexyl-TPP decreased the membrane potential to the same level as an equal concentration of FCCP, a strong uncoupler of oxidative phosphorylation (Figure 6a). Since the drop of the membrane potential has been proposed to cause the impairment of mitochondrial translation^39,40^ and is crucial for the import of mitochondrial protein precursors from the cytosol^41^; we further hypothesised, that the drop of the membrane potential could indeed be the main trigger of decreased IMM protein levels. However, 24-hours treatment with 2 μM FCCP did not result in the decrease of selected IMM steady-state RC protein levels compared to chronic treatment with hexyl-TPP, suggesting that the decrease in membrane potential alone is not causative of the effect of hexyl-TPP (Figure 6B, *S14*). This result was further confirmed when the drug exposure was extended to 48 hours. While the decrease of steady-state protein levels after 2 μM hexyl-TPP treatment was even more pronounced, the steady state protein levels in cells treated with FCCP remained unchanged (Figure S8A-B, *S18F*).

**Figure 6.**
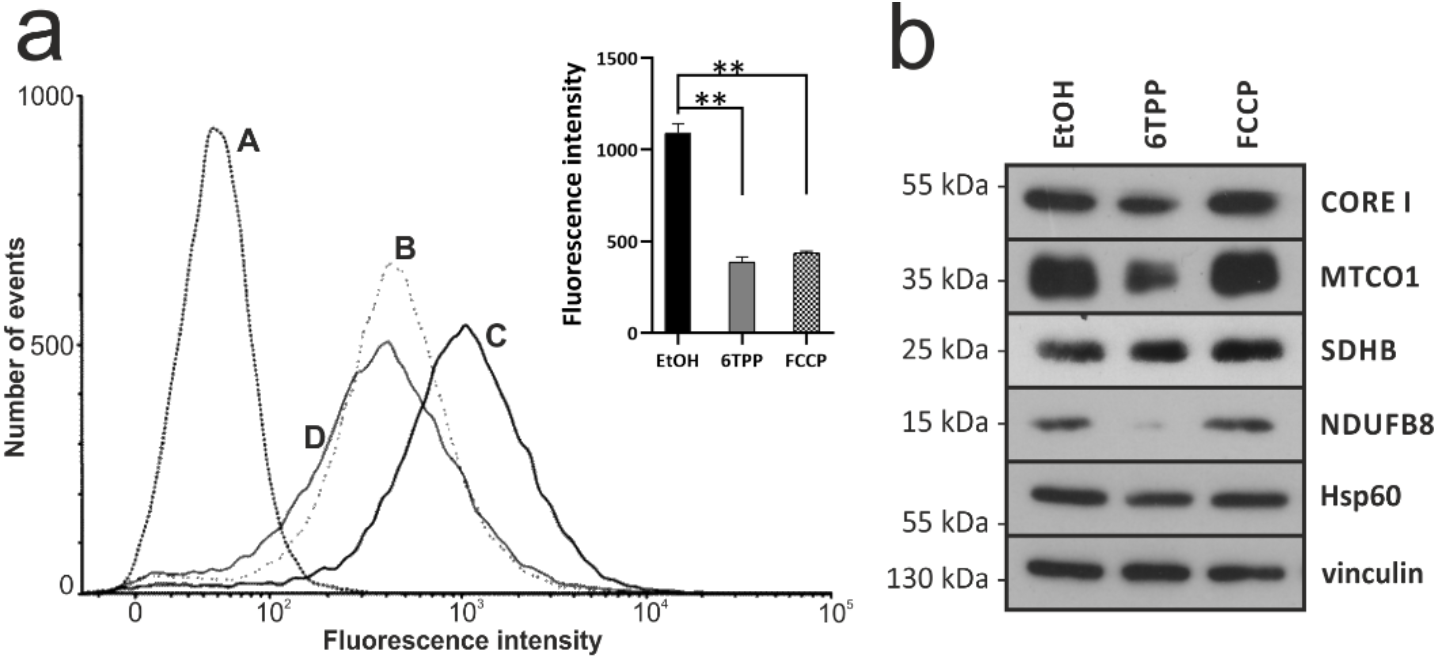
Drop in membrane potential alone is not causative of the effect of hexyl-TPP. **A**. The effect of 24 hours treatment of 2 μM hexyl-TPP or 2 μM FCCP compared to control treated with vehicle (EtOH) on mitochondrial membrane potential in C2C12 cells. Cells were stained with 50 nM TMRM, followed by flow cytometry analysis. A: unstained EtOH, B: 2 μM FCCP C: EtOH D: 2 μM hexyl-TPP. The geometric mean fluorescence intensities are shown in a representative histogram. The average of three independent experiments, is shown in a bar chart, error bars indicate mean ± SEM, ** indicate p < 0.01 in unpaired student t-test. **B.** Steady-state protein levels of different OXPHOS subunits from C2C12 cells treated with vehicle (EtOH) or 2 μM FCCP/2 μM hexyl-TPP for 24 hours as detected by immunoblotting. Vinculin and Hsp60 were used as loading controls. Scan of original immunoblot can be found in Fig. S14.

### 5. The effects of hexyl-TPP on mitochondrial structure and function are reversible

Previous studies have shown that TPP derivatives disrupt mitochondrial bioenergetics^19,21^ and here we show that these functional changes are accompanied by changes in mitochondrial morphology as well as a selective reduction of IMM proteins. We therefore investigated if both these effects could be reversed (and how quickly) when the offending chemical was removed.

Cells were treated with hexyl-TPP for 24 hours, and then incubated in fresh medium without TPP for 1 or 24 hours (Figure 7A-D). In all the rescue experiments, the hexyl-TPP medium was replaced twice, in a 30 minutes interval, to remove the TPP released from cells^10^. After just one hour of removing the compound, we observed that mitochondria were less swollen, although the mitochondrial network was still fragmented. After 24 hours without TPP, cells completely recovered their mitochondrial network to a state comparable to control cells. The functional disruptions in mitochondrial respiration also returned to control levels after 24 hours of new media incubation (Figure 7E). Finally, we assessed the possibility of a recovery of the mitochondrial protein levels after the removal of hexyl-TPP at 24-, 48- and 72-hours. Whilst protein levels were still significantly reduced 24 hours after the removal of TPP, we observed a gradual increase in protein levels after 48 and 72 hours (Figure 7F, Fig. S9, *S15*). Together, these data indicate that the effects of TPP derivatives on mitochondrial morphology, bioenergetics and IMM protein levels are reversible over time, when the TPP derivative is removed.

**Figure 7.**
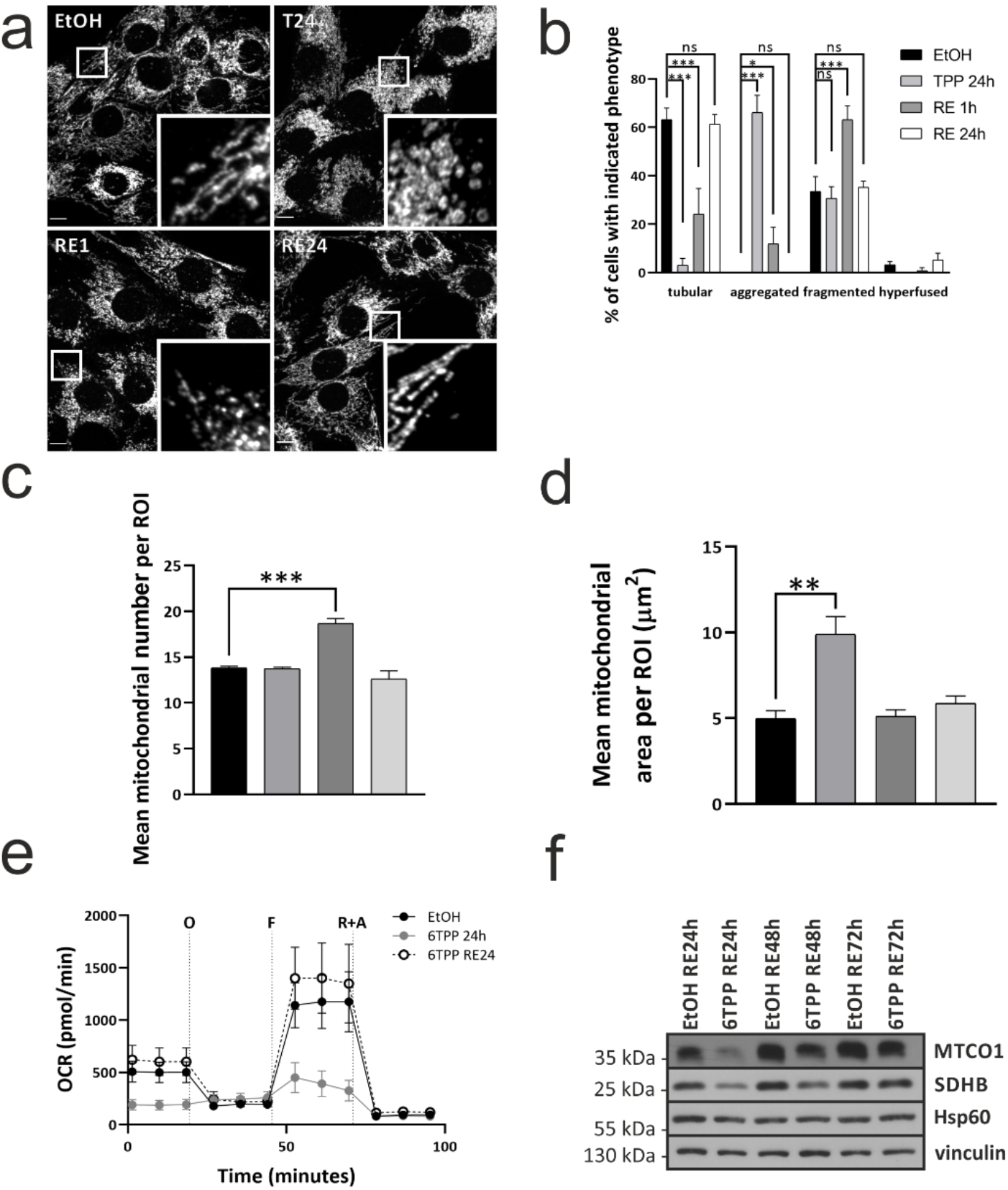
The effect of chronic treatment with hexyl-TPP is fully reversible. **A**. Representative confocal images of mitochondrial morphology in C2C12 cells treated with vehicle (EtOH) or 2 μM hexyl-TPP for 24 hours, followed by washes and incubation in cell growth medium without the treatment for 1 or 24 hours. Mitochondria were labelled using an anti-TOM20 antibody. Scale bars, 10 μm. **B.** Quantification of mitochondrial morphology from (A) showing three independent experiments, for each at least 40 cells have been quantified per condition, error bars are presented as mean ± SEM, n = 3, two-way ANOVA and Tukey’s multiple-comparisons test was used. **C.** Mitochondrial morphology quantified for mean mitochondrial number per ROI. **D.** Mitochondrial morphology quantified for mean mitochondrial area per mitochondrion. All data for C and D are shown as mean ± SEM, at least 15 cells for each experiment were quantified; three independent experiments were performed. Ordinary one-way ANOVA and Tukey’s multiple comparisons test was used. **E**. Seahorse analysis of oxygen consumption rate in C2C12 cells from vehicle (EtOH) compared to hexyl-TPP treatment for 24 hours and 24 hours treatment with hexyl-TPP followed by 24-hour incubation in cell growth medium without the treatment. Cells were seeded in a Seahorse XF-24 plate and incubated overnight. Cellular media was replaced with XF Assay Media and allowed to be equilibrate for 1h at 37 °C, CO_2_ free incubator. Basal respiration was measured, followed by sequential injections of 1 μM oligomycin (O), 2 μM FCCP (F) and 1 μM rotenone + 1 μM antimycin (R+A). All data are mean ± SEM, n=3. **F.** C2C12 cells were treated with vehicle (EtOH) or 2 μM hexyl-TPP for 24 hours, followed by washes and incubation in cell growth medium without the treatment for 24, 48 and 72 hours, the steady state protein levels of rescue experiment were detected on immunoblot. Vinculin and Hsp60 were used as loading controls.

### 6. Similar effects of the TPP moiety can be observed in MitoQ but not in the less hydrophobic TPP-hexanol

To understand if the effects described in this study were specific to alkyl-TPP derivatives or can be observed in other TPP compounds, we tested two different TPP derivatives, namely the mitochondrially targeted antioxidant MitoQ and a hydrophilic analogue of hexyl-TPP, TPP-hexanol (Fig. S1D-E).

Treatment with MitoQ led to a decrease in the steady state protein levels of IMM proteins even at a tenfold lower concentration compared to hexyl-TPP (Figure 8A, *S16*). This result likely reflects the differences in overall hydrophobicity between the two molecules with MitoQ being more hydrophobic. On the contrary, the treatment with TPP-hexanol did not result in a decrease in the selected steady state protein levels even at concentrations as high as 100 μM (Figure 8B, *S17*), most likely due to the substantially lower hydrophobicity caused by the hydroxyl group.

**Figure 8.**
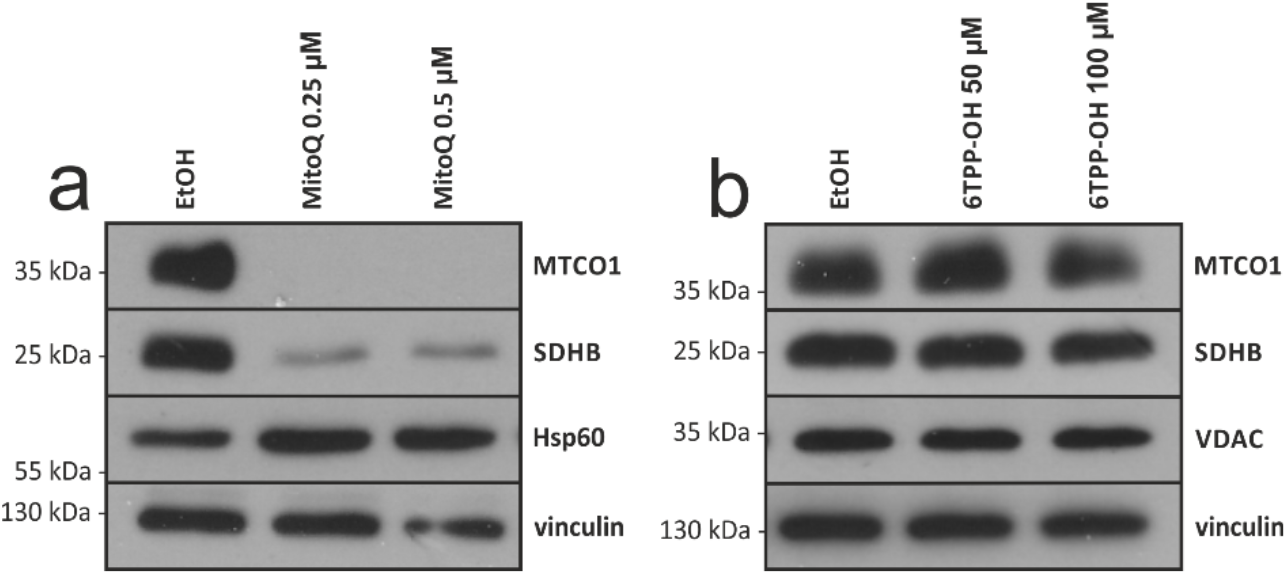
Effects of TPP derivatives vary with varying hydrophobicity. **A.** Immunoblot analysis for steady-state levels of respiratory chain complex IV (anti-MTCO1) and complex II (anti-SDHB) from C2C12 cells treated with vehicle (EtOH) or different concentrations of MitoQ for 24h. **B**. Immunoblot analysis for steady-state levels of respiratory chain complex IV (anti-MTCO1) and complex II (anti-SDHB) from C2C12 cells treated with vehicle (EtOH) or different concentration of hexanol-TPP. Vinculin and Hsp60 were used as loading controls.

## DISCUSSION

Lipophilic cations based on the triphenylphosphonium moiety used to target molecules into mitochondria are well known to associate with phospholipid biomembranes, particularly the inner mitochondrial membrane^42^ and have been shown to interact directly with cellular proteins, in particular Hsp70^43^. We therefore hypothesised that in addition to disrupting mitochondrial bioenergetics they may also disrupt specific proteins, such as subunits of the respiratory chain complexes and/or cause a disruption of the assembly of its supercomplexes. In the experiments presented here we observed that exposure of cells to alkyl-TPP derivatives decreases the steady-state protein levels of different subunits of respiratory chain complexes (MTCO1, NDUFB8, SDHB) in a concentration- and time-dependent manner (Figure 1A-C, 2A, Fig.S2A-C and Fig.S3A) and also with increasing potency as the hydrophobicity increases. Interestingly, the decrease of supercomplexes as well as steady-state protein levels was observed prior to the decrease in the abundance of ‘free’ complexes (Figure 2A-C, Fig.S3A). We also show that earlier steps of mitochondrial protein expression, i.e. mtDNA levels and transcription, are not negatively impacted by a TPP treatment (Figure 2D). This result suggests that the treatment with hexyl-TPP has a rapid detrimental effect on IMM protein complexes, possibly due to the degradation of their components.

When we examined the effect of hexyl-TPP treatment on the levels of a range of different mitochondrial proteins, we found out that the proteins of the OMM and matrix did not decrease in the treated cells (Figure 3A), while some but not all the IMM and intermembrane space proteins were affected (Figure 3B-D). These results suggest that the mechanism of TPP action is rather selective in targeting proteins of the IMM. We observed a selective reduction in OXPHOS components and the MCUC machinery, which are responsible for mitochondrial respiration and the regulation of calcium entry into the mitochondrial matrix, respectively. This selective behaviour might be due to a possible link between these two functional systems. The levels of calcium inside mitochondria have been shown to directly impact the activity of the Krebs cycle enzymes^44^, activity of complex IV^45^ or ATP synthase^46^, and MCU itself directly interacts with ATP synthase^47^. The mitochondrial calcium uniporter regulator 1 (MCUR1) has also been described as a scaffold protein for both MCU^48^ and cytochrome c oxidase^49^. The observed decrease in MCUC protein levels therefore could also be a result of an impairment of OXPHOS complexes by the TPP compounds.

The mitochondrial membrane remodelling in response to stress, such as a drop of a membrane potential, is, in part, regulated by the IMM GTPase OPA1. Stress-induced OPA1 processing regulated by the IMM metallo-protease OMA1 causes an accumulation of its shorter forms, which results in mitochondrial fragmentation and cristae architecture remodelling^33^. The exposure to hexyl-TPP did indeed lead to OPA1 cleavage (Figure 3E, Fig. S5) suggesting that this regulator protein could be involved in the mitochondrial membrane remodelling events observed. However, the fact that a decrease of IMM is observed concurrently with OPA1 cleavage rather than after it and that an siRNA knockdown of the protease OMA1 did not decrease the effect of TPP on IMM proteins suggest that OMA1 activation may not be a key player in the TPP-induced degradation (Fig. S6), but rather be concomitant effect of the TPP exposure.

Previous studies have shown that a drop in the mitochondrial membrane potential results in a disruption of mitochondrial translation^39,40^ and that the membrane potential is crucial for the import of mitochondrial protein precursors from the cytosol^41^. Therefore, we hypothesized, that the drop of the mitochondrial membrane potential due to the treatment with hexyl-TPP (Figure 6, Fig. S8) might be a cause for the observed decrease in IMM protein levels. However, our results showed that an equal decrease in membrane potential caused by 2 μM of the uncoupler FCCP during equal times (Figure 6A), did not lead to the same decrease in steady state protein levels (Figure 6B, Fig. S8). This rules-out the possibility of the protein decrease being simply due to the dissipation of membrane potential and suggests a direct role for hexyl-TPP on mitochondrial protein synthesis and/or stability and/or degradation, by so far uncharacterised mechanisms. The specific effect of lipophilic TPP-based cations hints at a currently unknown process linking the disruption of the IMM to the changes in membrane protein levels, which warrants further study.

As previously reported for the more hydrophobic dodecyl-TPP^17^, we observed that exposing cells to 2 μM hexyl-TPP causes significant changes in mitochondrial morphology, including mitochondrial fragmentation and swelling (Figure 4A-F, Fig.S7A-D). Using electron microscopy, we showed loss of cristae architecture but, perhaps surprisingly, electron tomography analysis did not find any ruptures of the IMM (Figure 5A-G). Such a result goes against the often-believed hypothesis that TPP derivatives behave as detergents virtually “dissolving” the IMM. The hydrophobicity of the TPP-linked molecules is clearly the most relevant physico-chemical parameter. This phenomenon is evident since the more hydrophobic mitochondria-targeted antioxidant MitoQ exhibits stronger effects on IMM protein levels compared to the less hydrophobic congener of hexyl-TPP, TPP-hexanol (Figure 8A-B).

Finally, we proposed that alterations in the mitochondrial function caused by hexyl-TPP treatment are fully reversible upon the removal of the molecule (Figure 7A-F, Fig. S9). This finding further strengthens the hypothesis that TPP-induced changes are the result of controlled processes, not due to non-specific toxicity.

The results presented are relevant both for the design of new mitochondrial probes and in the development of new mitochondria-targeted drugs. This study also provides a novel piece into the puzzle of interpreting the effect of TPP-conjugated molecules. When new such compounds are created and tested the effects of the supposedly biologically active payload (be it antioxidant, cytotoxic agent, hormone mimetic, ETC inhibitor or any other interesting molecule) should be always compared to the effects of a an “inert” carrier, i.e. a TPP-based compound of similar physical-chemical properties (mainly hydrophobicity) but without the biologically active moiety as a “negative” control. Without such controls the observed effects may be fully due to the lipophilic cationic nature of the compound and the addition of a pharmacologically active payload may have been unnecessary.

Our experiments looked at the effects of the TPP moiety after a relatively long *in vitro* exposure. *In vivo*, it is conceivable that the excretion and redistribution of the compound among organs and tissues would result in a fluctuation between the disrupted and recovery phenotypes we described in this study. On the other hand, in certain cells/tissues or under certain circumstances this might not be the case and these *in vitro* effects might be observed also *in vivo;* e.g. in the kidney, where the compounds may be concentrated over time, due to being the main route of excretion^50^. Importantly, there are already first descriptions of the negative effects of TPP treatment in different tissues *in vivo* models^51^.

The present study goes well beyond the previously published data on the relationship of chain length/hydrophobicity of TPP compounds on their toxicity/cellular effects. The significance of our paper lies in the fact that we show for the first time a specific and reversible effect of alkyl-TPP compounds (i.e. supposedly “inert” targeting moieties without a biologically active payload) on inner mitochondrial membrane proteins, which correlates well with previously seen functional effects and effects on supercomplex levels shown by Rohlenova et al.^11^. We further test (and refute) the mechanistic hypotheses 1) that the loss of protein is caused primarily by the loss of IMM potential (this is the reason for comparing the effects of FCCP and alkyl-TPP and measuring the membrane potential) and 2) that it is primarily mediated by OMA1 activation. As such our study significantly furthers our understanding of how lipophilic cations with or without any biologically active compounds interact with the inner mitochondrial membrane.

Our paper is novel in that it shows that the TPP moiety itself decreases steady-state protein levels of IMM, disturbs the native structure of OXPHOS complexes, and results in significant morphological changes and mitochondrial swelling but does not seem to cause the ruptures in IMM. Indeed, some of these were shown before, however, always related to antioxidants or drugs conjugated to TPP molecules. However as far as we know, there is no paper showing that some of these disruptive effects are related to TPP moiety itself.

We can conclude here that the TPP moiety used for the synthesis of a range of mitochondria-targeted compounds is not inert but initiates profound, specific and reversible effects on the mitochondrial structure and function, which must be taken into account whenever these compounds are used.

## ACKNOWLEDGEMENT

We would like to thank Dr. Erika Fernandez-Vizarra and Dr. Hiran Prag, both of whom provided feedback on an earlier version of the manuscript and to Zuzana Varaliova for performing qPCR to test siRNA efficiency. Any errors in this work are the author’s own and should not reflect in any way on the contributions from these valued supporters.

## AUTHOR CONTRIBUTIONS

JT and AK conceived the study; AK, FT, VŠ, MV, TB performed experiments and AK, LT, MV, TB analysed data; JT, JP, JN contributed to methods; AK and JT wrote the manuscript, all co-authors participated in reviewing the manuscript.

## FUNDING

This work was supported by PROGRES-Q36-METAB-GAP UK from Charles University, and by the Medical Research Council, UK (MC_ UU_00015/7) for JP. The electron microscopy (Biology Centre, České Budějovice) analyses were supported by the MEYS CR (LM2018129 Czech-BioImaging) and ERD Fund (003/0000441 and CZ.02.1.01/0.0/0.0/16_013/0001775). LT is an MRC-funded PhD student.

## COMPETING INTERESTS

The authors declare no competing interests.

