## Supplementary figures and data for "Selective and reversible disruption of mitochondrial inner membrane protein complexes by lipophilic cations"

**This supplementary material includes:**

Figures S1 to S14.

Figures S6 – S14 show full scans of the immunoblots, or the original membranes of any immunoblot shown in the main text of the paper. The cropped bands are framed and any membrane exposure which was not made at the same time is specifically labelled. The membranes developed on a digital Amersham Imager 680 are shown as jpg images which allows to present them together with the marker (Thermo Fisher Scientific, 26619). For processing of the main figures in Fiji the equivalent tiff files of these images were used, these are available from the corresponding author on request, similarly as any other raw data from the paper.

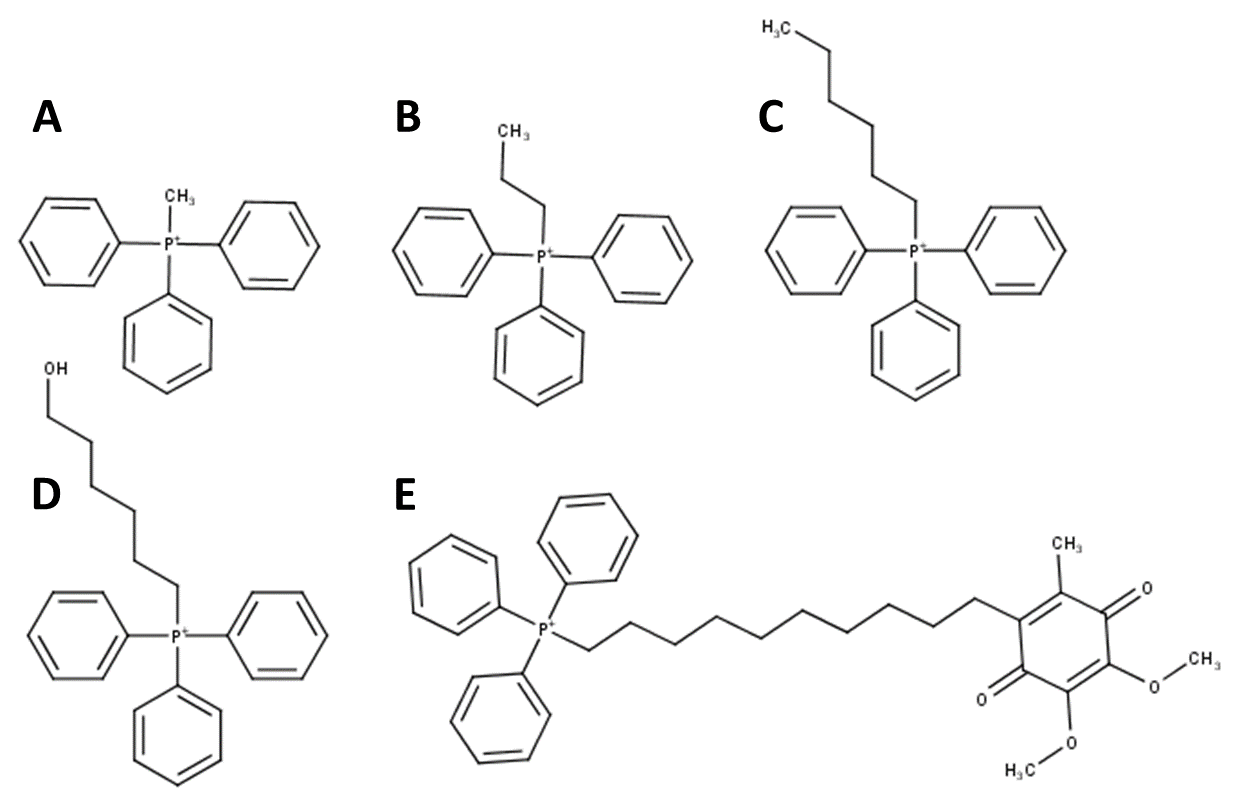

***Fig. S1 Structures of TPP derivatives used within the study.*** ***A****. methyltriphenylphosphonium* ***B.*** *propyltriphenylphosphonium* ***C****. hexyltriphenylphosphonium* ***D.*** *triphenylphosphonium hexanol* ***E.*** *MitoQ*

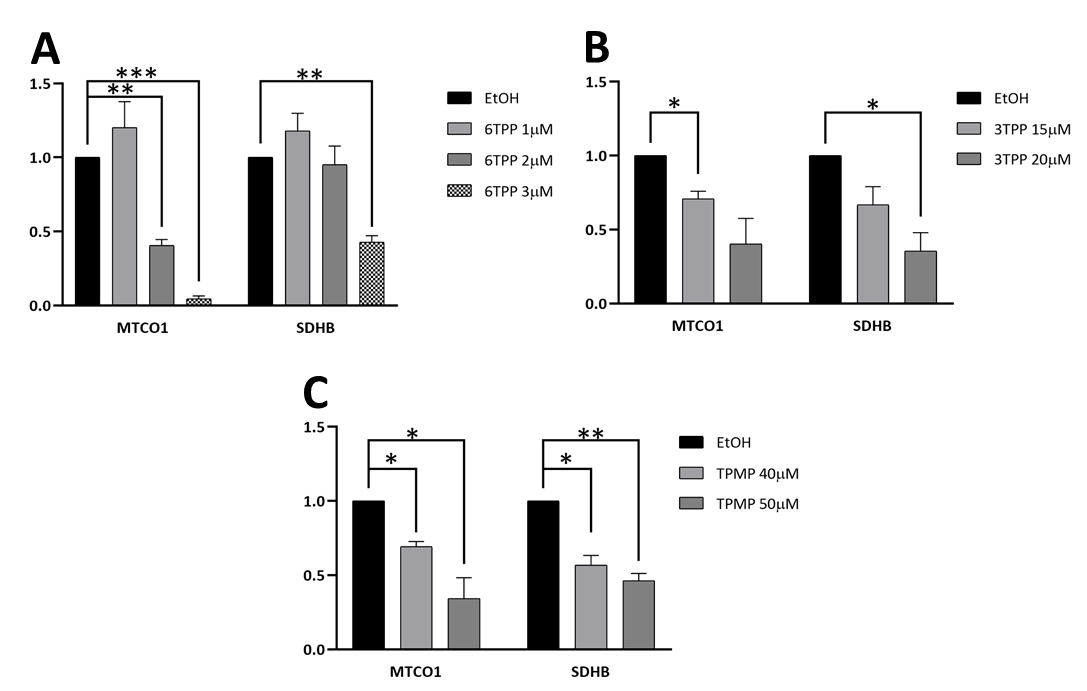

***Fig. S2. Treatment with TPP derivatives leads to a decrease in selected subunits of OXPHOS complexes.*** *The graphs A-C show the densitometric quantification from Figure 1 a-c respectively, the signal corresponds to subunits of complex IV (MTCO1) and complex II (SDHB) and is normalized to Hsp60. The mean of the three control samples was set to 1 and all the measurements were referenced to that value. Error bars are presented as ± SEM, * indicate p < 0.05, **p < 0.01, ***p < 0.001 in unpaired student t-test.*

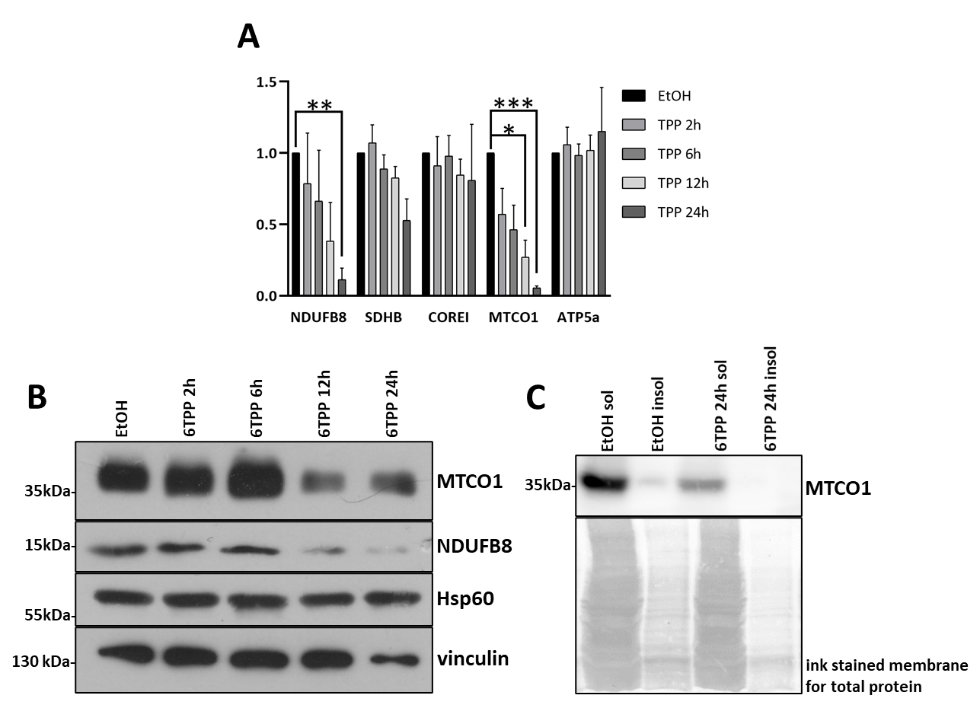

**Fig. S3 Treatment with hexyl-TPP leads to a decrease of OXPHOS complexes in a time dependent manner in different cell lines A.** The graph shows the densitometric quantification from Figure 2a, the signal corresponds to subunits of complex I (NDUFB8), complex II (SDHB), complex III (COREI), complex IV (MTCO1) and ATP synthase (ATP5a) and is normalized to Hsp60. The mean of the three control samples was set to 1 and all the measurements were referenced to that value. Error bars are presented as ± SEM, * indicate p < 0.05, **p < 0.01, ***p < 0.001 in unpaired student t-test. **B.** Steady-state levels of OxPhos components from HepG2 cells treated with vehicle or 2 µM hexyl-TPP over different timepoints. Vinculin and Hsp60 were used as loading controls. Scan of original immunoblot can be found in fig. S14 A. **C**. Mild cellular lysis followed by SDS extraction of insoluble pelleted proteins detected on immunoblot; sol (soluble) insol (insoluble) fraction.

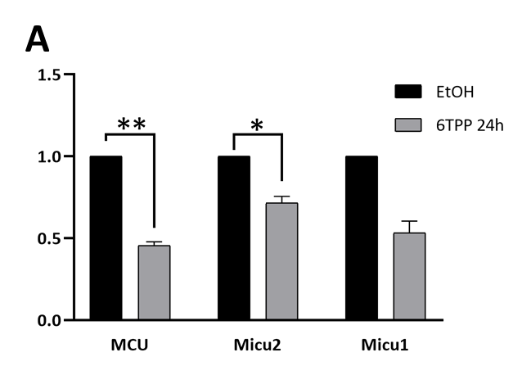

***Fig. S4 Treatment with hexyl-TPP leads to the decrease of MCUC subunits A.*** *The graph shows the densitometric quantification from Figure 3d, the signal corresponds to MCUC subunits (MCU, Micu2 and Micu1) and is normalized to Grp75. The mean of the three control samples was set to 1 and all the measurements were referenced to that value. Error bars are presented as ± SEM, * indicate p < 0.05, **p < 0.01 in unpaired student t-test.*

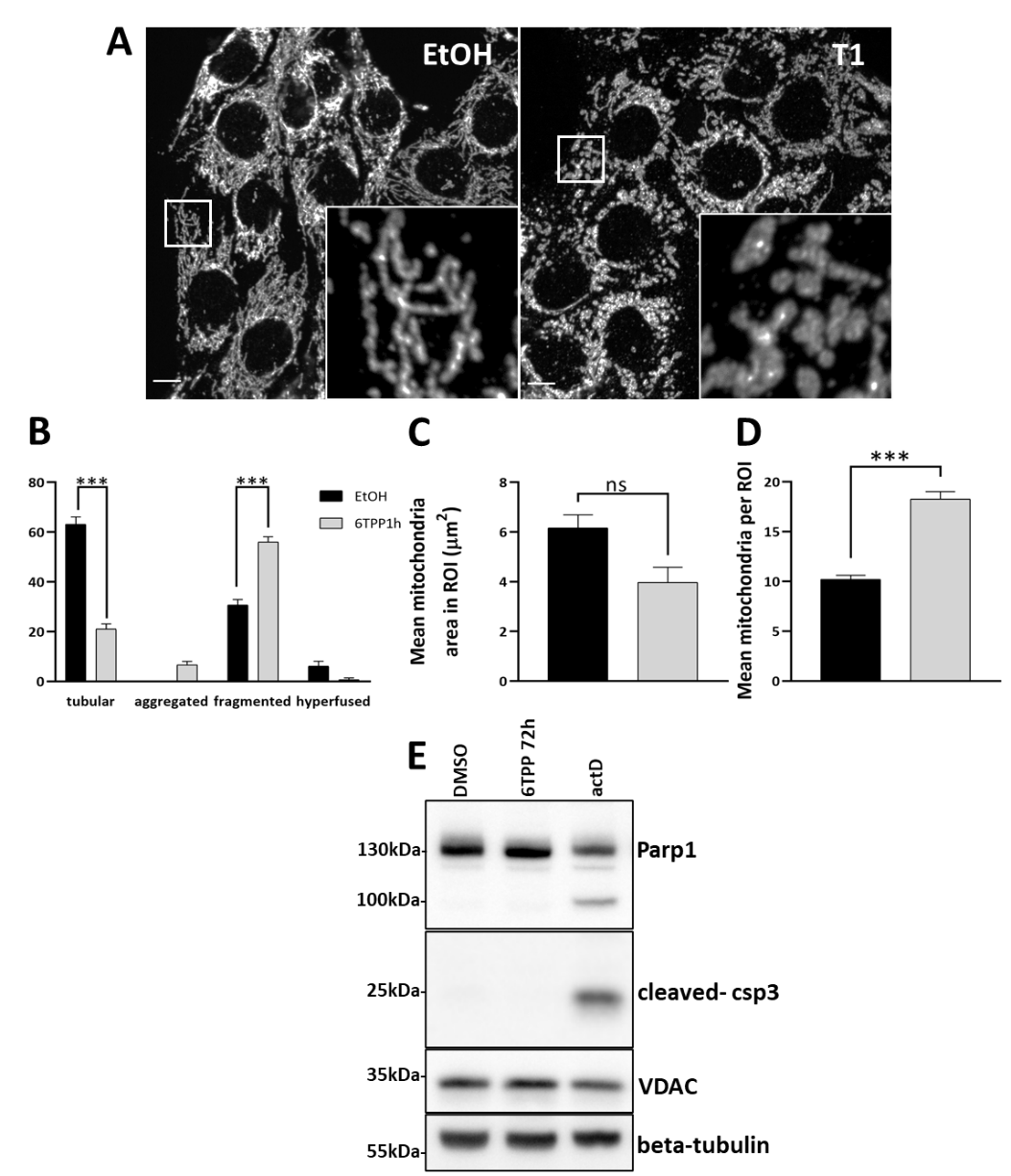

**Fig. S5 Treatment with hexyl-TPP alters mitochondrial morphology but does not induce apoptosis. A**. Representative images of mitochondrial morphology in C2C12 cells treated with vehicle (EtOH) or 2 µM hexyl-TPP for 1hour. Mitochondria were labelled using an anti-TOM20 antibody. Scale bars 10 µm. **B.** Quantification of mitochondrial morphology from A showing three independent experiments, for each at least 50 cells have been quantified, error bars are presented as mean ± SEM, n = 3, two-way ANOVA and Tukey´s multiple-comparisons test was used. **C.** Mitochondrial morphology quantified for mean mitochondrial number for each region of interest (ROI). **D.** Mitochondrial morphology quantified for mean mitochondrial area per mitochondrion. All data for C and D are shown as mean ± SEM, at least 15 cells for each experiment were quantified; three independent experiments were performed. Ordinary one-way ANOVA and Tukey´s multiple comparisons test was used. **E.** Representative immunoblot analysis for apoptosis-related proteins (cleaved caspase-3 and PARP1). C2C12 cells were exposed to vehicle or 2 µM hexyl-TPP for 72 hours, actinomycin D (actD) was used as a positive control. Scan of original immunoblot can be found in fig. S14 B.

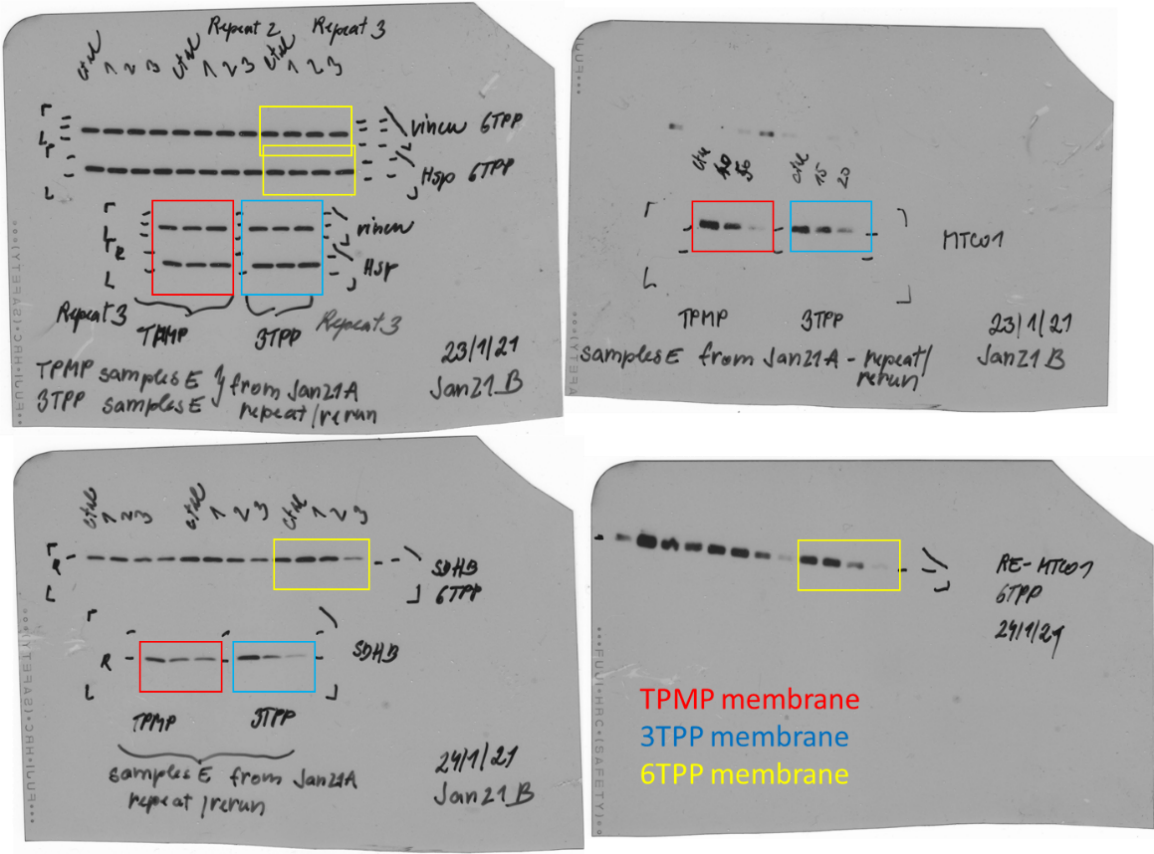

***Fig. S6 Full scans of Immunoblots from Fig. 1a, b, c.***

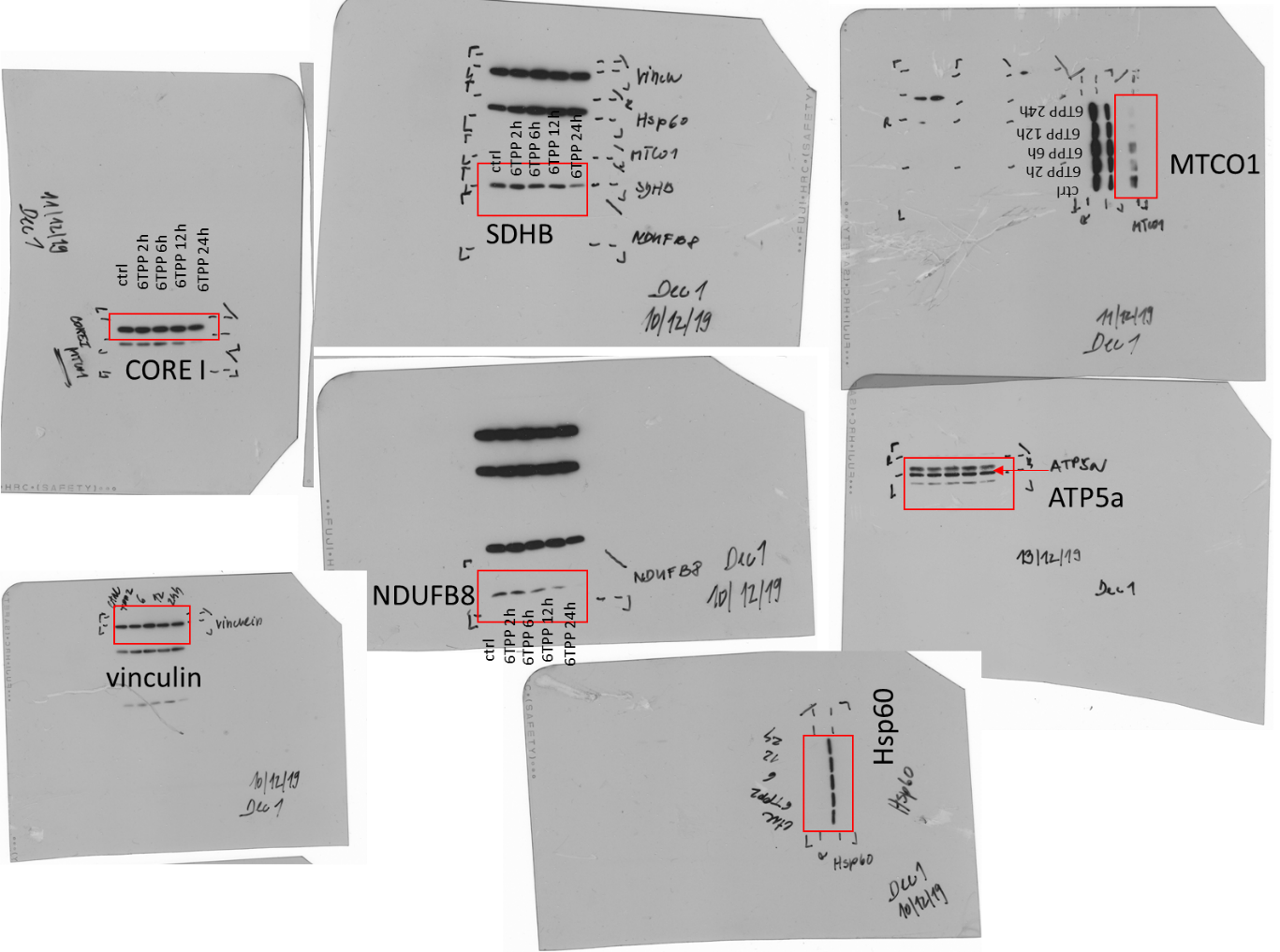

***Fig. S7 Full scans of Immunoblot from Fig. 2a.***

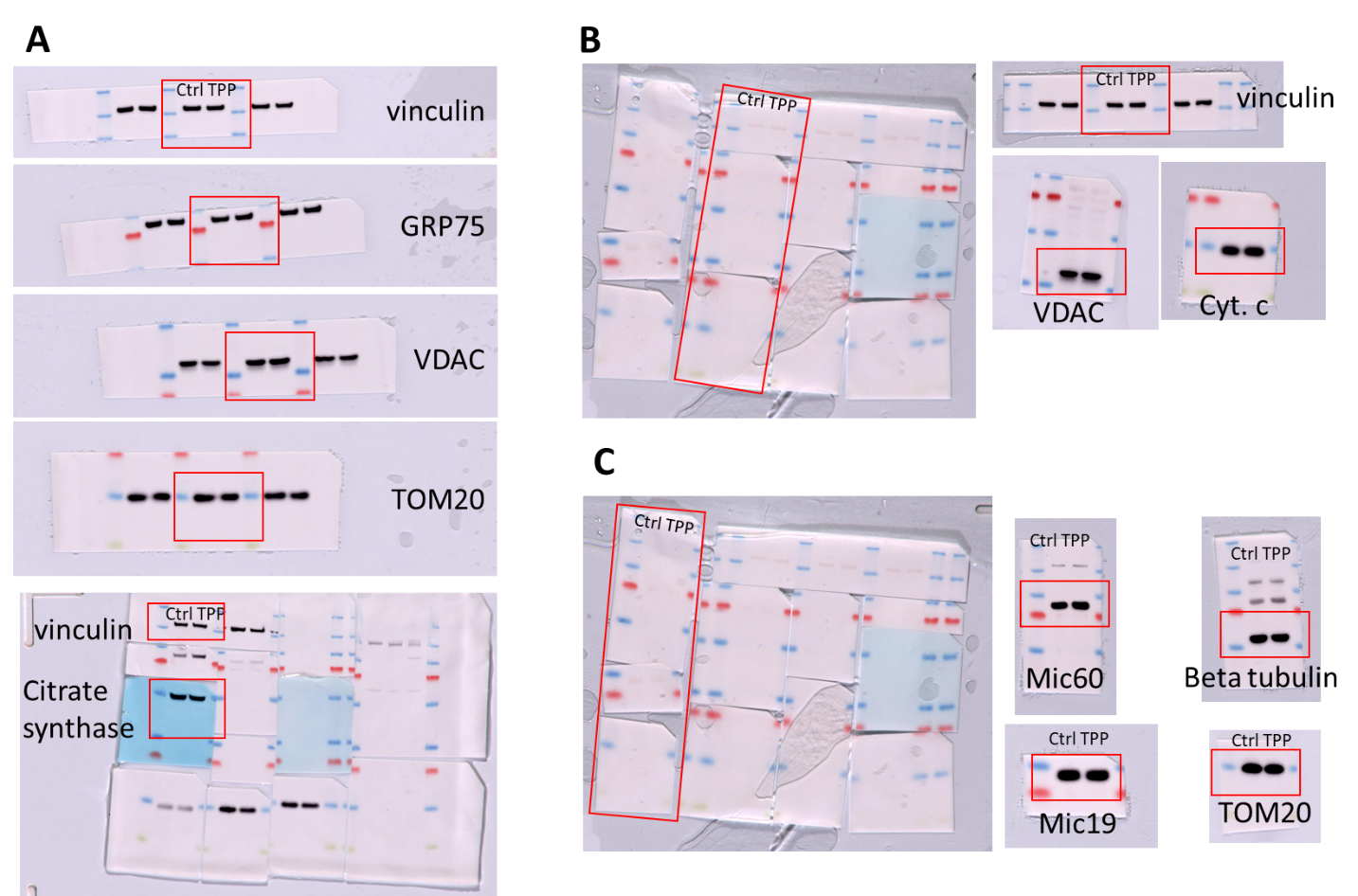

***Fig. S8 A, B, C Original membranes of Immunoblots from Fig. 3a – c, respectively.***

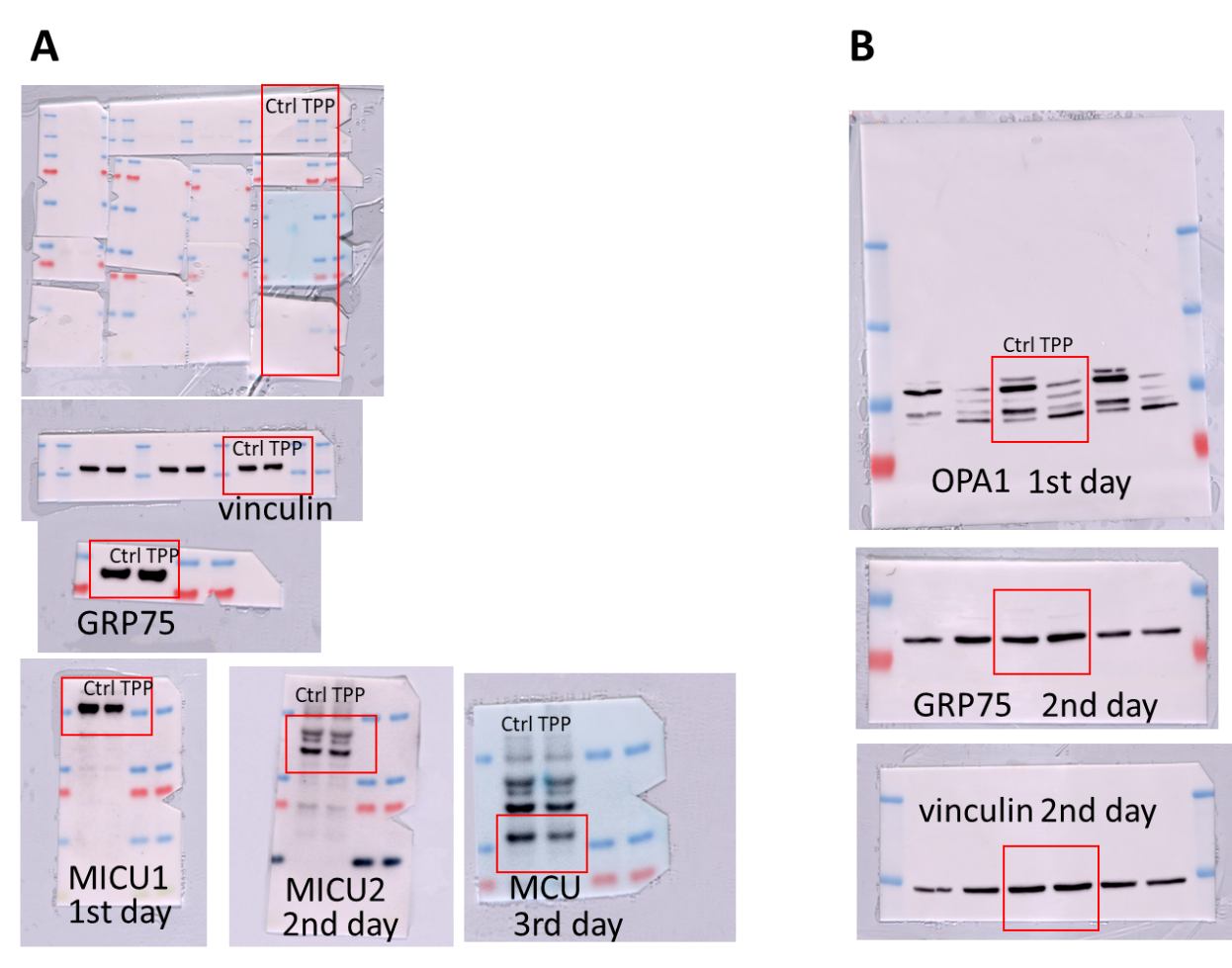

***Fig. S9 A, B Original membranes of Immunoblots from Fig. 3d – e, respectively.***

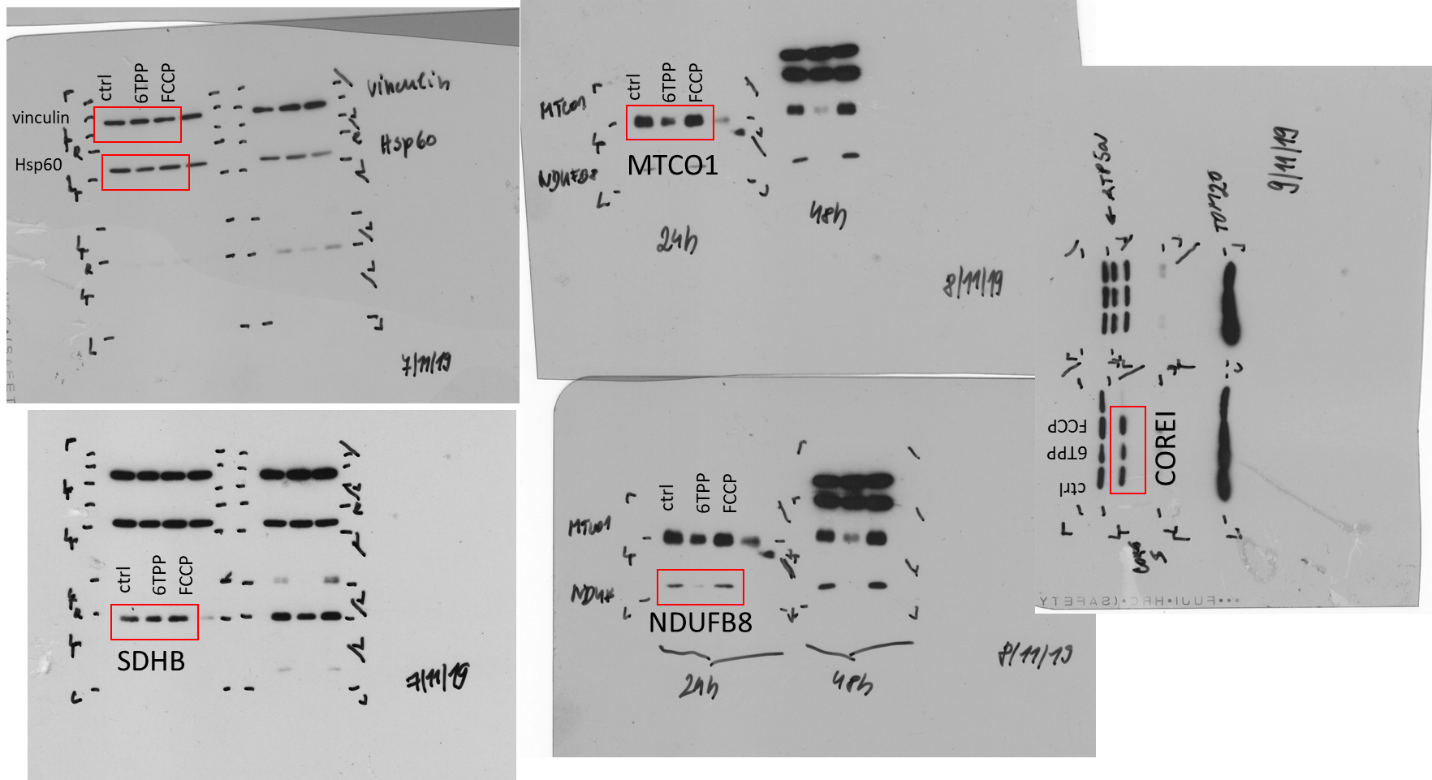

***Fig. S10 Full scans of Immunoblot from Fig. 6b.***

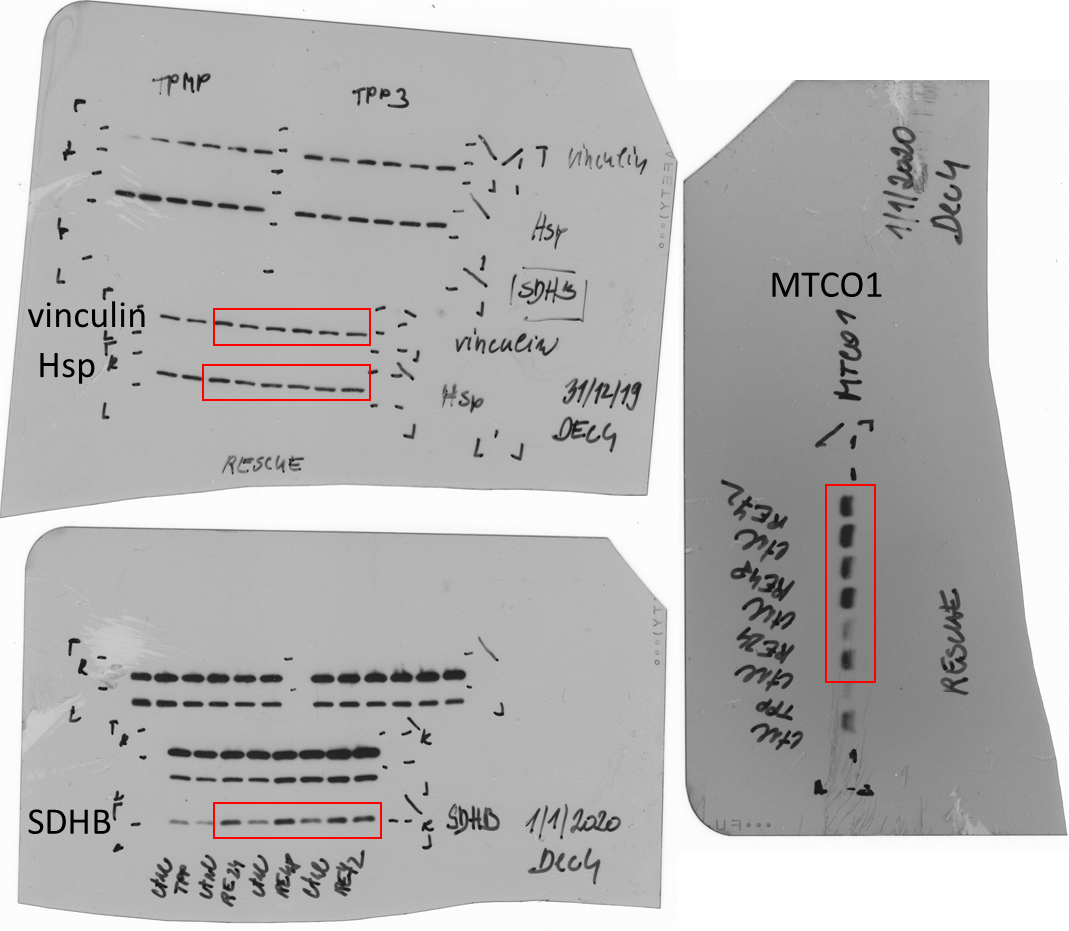

***Fig. S11 Full scans of Immunoblot from Fig. 7f.***

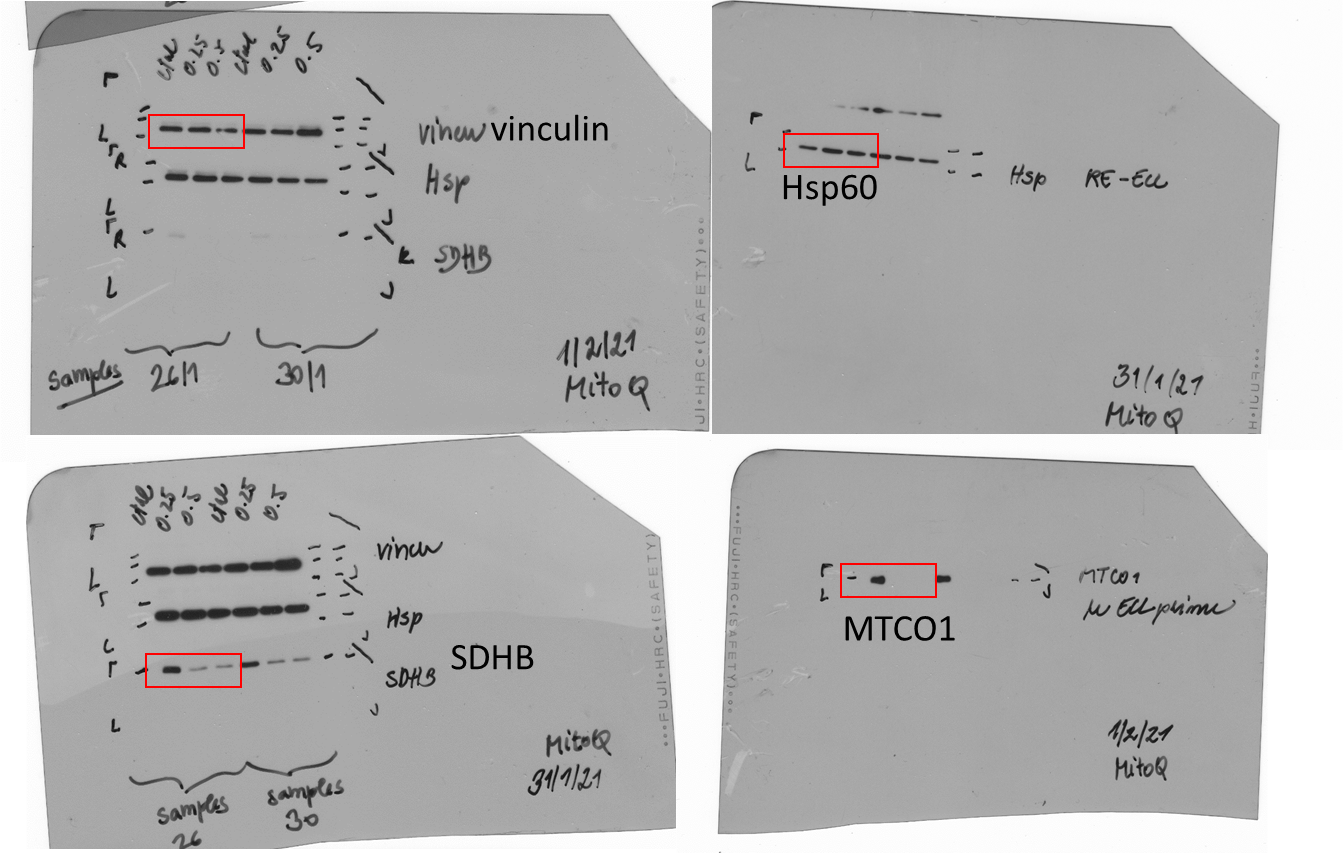

***Fig. S12 Full scans of Immunoblot from Fig. 8a.***

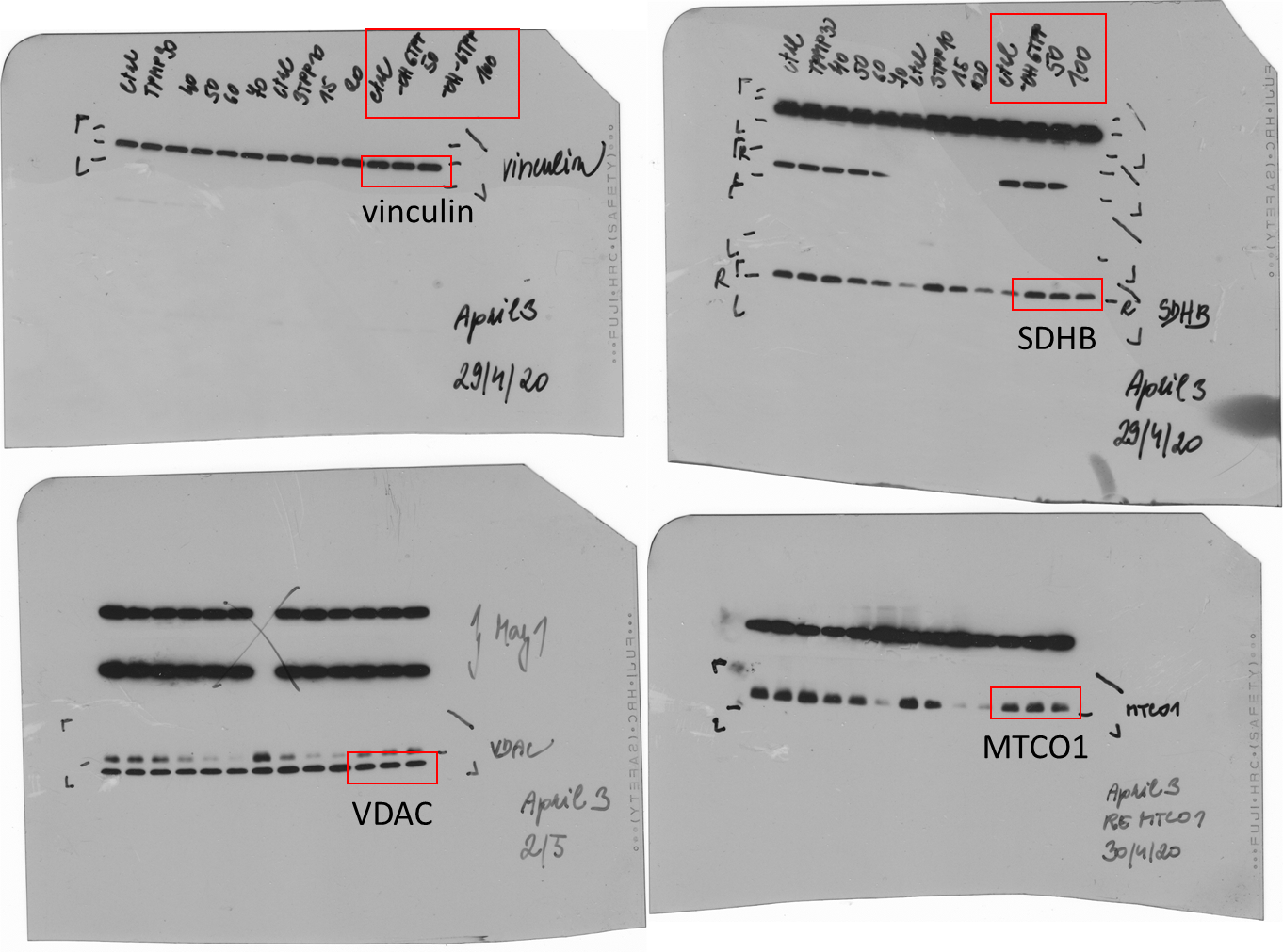

***Fig. S13 Full scans of Immunoblot from Fig. 8b.***

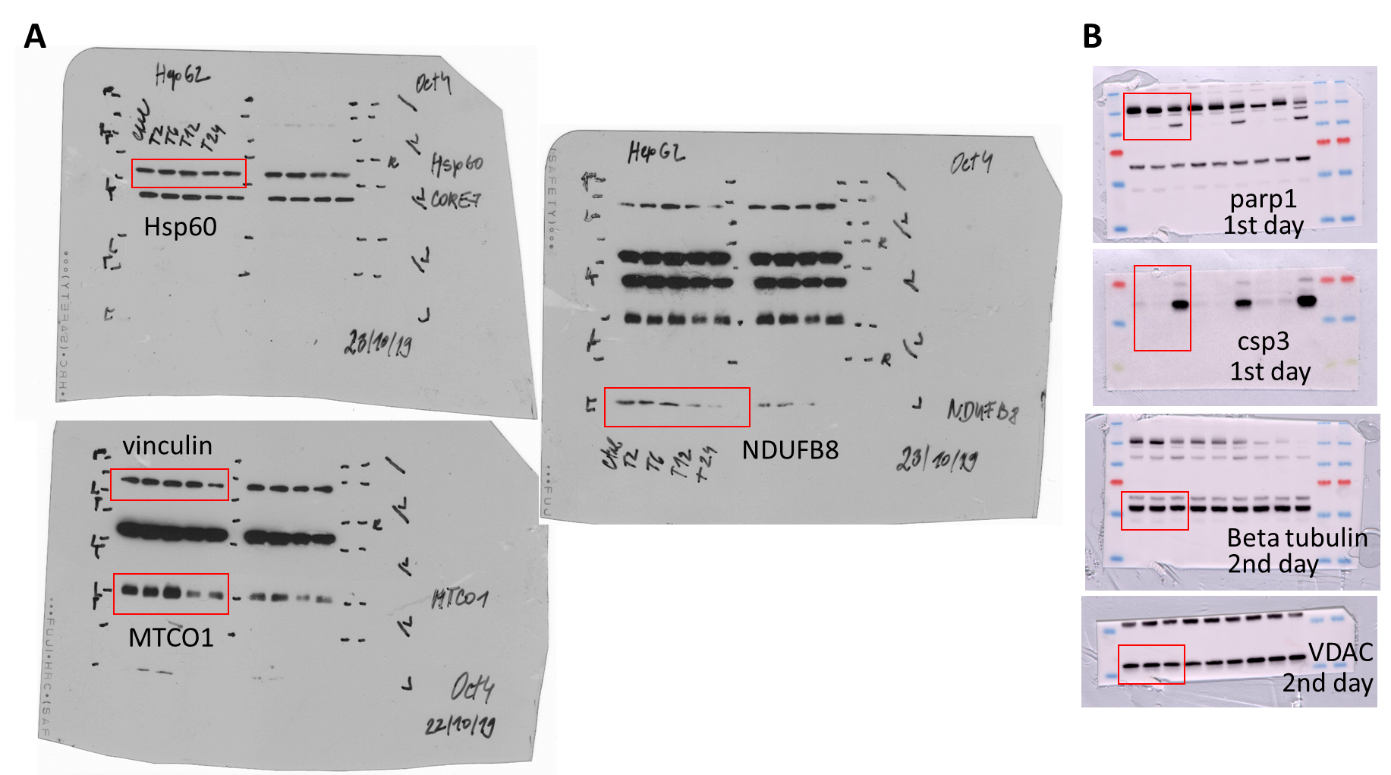

***Fig. S14 A Full scans of Immunoblots from fig. S3B. B Original membranes of Western Blots from fig. S5E.***
